# “When I talk about it, my eyes light up!” Impacts of a national laboratory internship on community college student success

**DOI:** 10.1101/2023.05.08.539924

**Authors:** Laleh E. Coté, Seth Van Doren, Astrid N. Zamora, Julio Jaramillo Salcido, Esther W. Law, Gabriel Otero Munoz, Aparna Manocha, Colette L. Flood, Anne M. Baranger

## Abstract

Participation in technical/research internships may improve undergraduate graduation rates and persistence in science, technology, engineering, and mathematics (STEM), yet little is known about the benefits of these activities a) for community college students, b) when hosted by national laboratories, and c) beyond the first few years after the internship. We applied Social Cognitive Career Theory (SCCT) to investigate alumni perspectives about how CCI at Lawrence Berkeley National Laboratory (LBNL) impacted their academic/career activities. We learned that alumni had low confidence and expectations of success in STEM as community college students. Participation in CCI increased their professional networks, expectations of success, and STEM skills, identity, and self-efficacy/confidence. Hispanic/Latinx alumni recalled the positive impact of mentors who prioritized personal connections, and women valued “warm” social environments. We propose several additions to the SCCT model, to better reflect the supports and barriers to STEM persistence for community college students.

## Introduction

National reports highlight the value of investing in science, technology, engineering, and mathematics (STEM) education to support the long-standing effort to increase the representation of women; Black, Hispanic/Latinx, and Native American people; and people with disabilities in these fields.^1–3^ One component of the recent CHIPS and Science Act is to take active steps toward broadening participation in STEM disciplines, with the ultimate goal of increasing diversity across the workforce.^4^ To promote diversity, equity, inclusion, and accessibility across their large suite of programs, the United States (U.S.) Department of Energy (DOE) Office of Science recently implemented the Reaching a New Energy Sciences Workforce (RENEW) initiative, the Promoting Inclusive and Equitable Research (PIER) component of research proposals, and other related efforts.^5^ These initiatives are important to higher education and workforce development, because STEM majors in the U.S. drop out of school or switch to a non-STEM major at higher rates than their peers in non-STEM disciplines, and this is even more likely for students who are Alaska Native, Black, Hispanic/Latinx, Native American, female, first-generation to college, and low-income.^6–12^ Thus, it is critical to explore those factors that contribute to student persistence in STEM and determine what encourages a student to “stay?”

Although student engagement in STEM research experiences and internships are an effective way to broaden participation and promote long-term retention in STEM, we know very little about these opportunities hosted at DOE national laboratories and/or those in which community college students participate. Additionally, there are few studies that examine the long-term perspectives of students after their participation in a technical/research experience.^13,14^ Thus, we have studied the experiences of community college STEM majors before, during, and after they participated in an internship at a DOE national laboratory and how they believe their experiences impacted their academic and career activities. We have investigated the following research questions:

RQ1: *Prior to applying to the Community College Internship (CCI) at Lawrence Berkeley National Laboratory (LBNL), what were the experiences of CCI alumni when they were community college students studying STEM?*
RQ2: *What skills, gains, and/or benefits do alumni of the CCI program at LBNL attribute to their participation in this program?*
RQ3: *In what ways do CCI alumni believe that their backgrounds, cultures, and identities impacted their experiences studying and pursuing careers in STEM?*

### Study Overview

We explored the ways in which community college students experience benefits as a direct result of participation in a STEM internship, to learn both how and why this program impacted its participants. Although many quantitative studies have documented achievement levels for undergraduates across different institution types, there is a gap in knowledge about the academic trajectories and experiences of students attending U.S. community colleges (Table 1) and the ways in which their personal experiences have impacted their academic and career success.^15,16^ Informed by many studies that link psychosocial, academic, and professional benefits for students who complete STEM research experiences and internships^17–19^; the lack of knowledge about the experiences of community college STEM majors; and the absence of studies documenting programs at DOE national laboratories, we considered the role of the CCI program on our study participants’ academic and career activities. We gained a deep understanding of participant experiences before, during, and after participation in the program through the documentation of their experiences as students and their opinions about the factors that influenced their academic/career trajectories (including their own community, culture, and background).

**Table 1.**
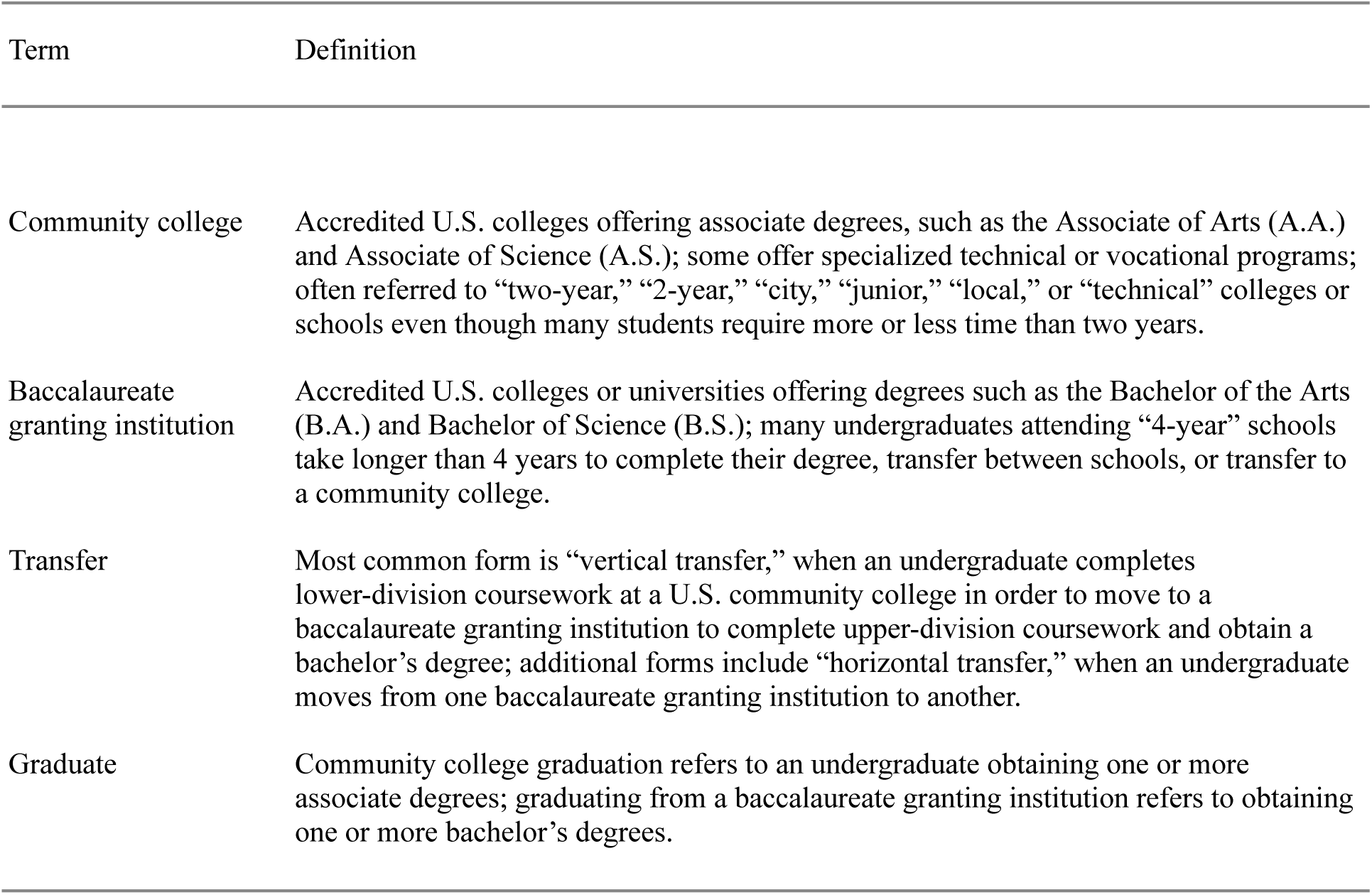
Definition of Terms.

| Term | Definition |
| --- | --- |
| Community college | Accredited U.S. colleges offering associate degrees, such as the Associate of Arts (A.A.) and Associate of Science (A.S.); some offer specialized technical or vocational programs; often referred to “two-year,” “2-year,” “city,” “junior,” “local,” or “technical” colleges or schools even though many students require more or less time than two years. |
| Baccalaureate granting institution | Accredited U.S. colleges or universities offering degrees such as the Bachelor of the Arts (B.A.) and Bachelor of Science (B.S.); many undergraduates attending “4-year” schools take longer than 4 years to complete their degree, transfer between schools, or transfer to a community college. |
| Transfer | Most common form is “vertical transfer,” when an undergraduate completes lower-division coursework at a U.S. community college in order to move to a baccalaureate granting institution to complete upper-division coursework and obtain a bachelor’s degree; additional forms include “horizontal transfer,” when an undergraduate moves from one baccalaureate granting institution to another. |
| Graduate | Community college graduation refers to an undergraduate obtaining one or more associate degrees; graduating from a baccalaureate granting institution refers to obtaining one or more bachelor’s degrees. |

### Asset-based approach

We take an **asset-based (anti-deficit) approach** to studying the experiences of individuals who began their STEM coursework as community college students. Deficit thinking involves labeling certain students as being “disadvantaged” or “lacking” in some way, and can be used to justify why some students “fail to achieve” at the same levels as students in other groups.^20,21^ A deficit-oriented approach might place responsibility on students for “leaving” STEM – because they are less prepared or motivated – as opposed to considering the ways in which institutions may be differentially serving students from different groups.^22–24^ In this study, we ask questions that allow us to investigate the reasons why community college students *persist* in STEM, aligned with the asset-based approaches taken by other educational research studies.^25–27^ If we identify the factors that have a positive influence on students, we better position ourselves to reproduce these supports in the future.

### Technical and research experiences benefit STEM majors at baccalaureate granting institutions

For students at baccalaureate granting institutions, numerous studies show that participation in a mentored research experience has many potential benefits, including increased academic achievement, likelihood of completing a STEM undergraduate degree, interest in completing a STEM graduate degree, and persistence in STEM.^2,18,28–32^ Working on technical projects, engaging in research, and receiving support from mentors can clarify students’ academic/career goals, and lead to gains in self-efficacy, confidence, technical skill level, and persistence in their field of study.^19,30,33^ Factors such as self-efficacy, STEM identity, and internalization of the values of the scientific community are thought to act as mediators between STEM activities and overall persistence in STEM careers.^34,35^

Undergraduates are greatly influenced by the activities they engage in during the first two years of their college experience, but many undergraduates do not participate in STEM technical work or research until the last two years of their bachelor’s degree.^36,37^ Studies suggest that participating in research during the first two years of undergraduate studies has the potential to increase student grade point average (GPA) at graduation, discipline-specific content knowledge, confidence, curiosity, interest, science identity, and institutional satisfaction.^38–40^ Aptly stated by Hagedorn and Purnamasari^41^, a student “will not elect to be a nuclear physicist” without some exposure to the field, or knowledge about what the career path entails; participation in research experiences or internships can provide these opportunities.

### STEM technical and research experiences for community college students

Collectively, many community college students are interested in transferring to baccalaureate granting institutions, graduating with a STEM degree, and entering the STEM workforce, but very few meet these goals.^42^ Of those “entering” students taking STEM coursework at a community college, 75-80% aspire to graduate with a bachelor’s degree, but only 15-16% of community college STEM majors achieve this goal.^43–47^ The President’s Council of Advisors on Science and Technology (PCAST) has called for DOE national laboratories to engage with community colleges in preparing a diverse future STEM workforce through internships and work-based learning experiences.^48^ This is, in part, due to the fact that community college students are collectively more diverse than students attending public or private baccalaureate granting institutions, with respect to gender, race, ethnicity, neurodiversity, disability, career pathway, parental educational attainment, and socio-economic status.^49–53^ Low levels of psychosocial constructs (e.g., STEM identity, confidence) are thought to be important to understanding the low graduation rates and STEM persistence of Black and Hispanic/Latinx undergraduates.^54^ Previous work has shown that undergraduate research and internships for undergraduates attending baccalaureate granting institutions can increase these psychosocial constructs, and internship programs hosted by DOE national laboratories are a possible mechanism through which community college students might similarly benefit.^19,33^

It is clear that engagement of community college students in STEM research experiences and internships could have enormous impact on their retention in STEM. However, although nearly 50% of people with STEM bachelor’s or master’s degrees attended a U.S. community college, an estimated 6% (or fewer) of the studies published each year about STEM research experiences and internships include data from community college or transfer students.^55–63^ This may be because fewer community college students participate in STEM research experiences and internships due to lack of access. Additionally, there may be a connection between concealable identities that “carry negative stereotypes” in the academic science community – such as attending a community college – and the lack of scholarly work dedicated to understanding those identities.^64^ There are some examples of programs designed for community college students that report positive outcomes^31,65–68^, but these are infrequently reported in the literature.^56^

### National laboratories’ role in STEM education is underrepresented in the literature

In the past few decades, some technical and meeting reports, conference papers, and abstracts have been published about STEM education and outreach activities at DOE national laboratories, and a small number of peer-reviewed publications on the subject.^69–76^ Beyond this, educational scholars at baccalaureate granting institutions – who produce the majority of studies about internships and research experiences for STEM students – rarely mention educational opportunities at national laboratories in their work. With national laboratories spending over $500 million annually to provide students, postdoctoral scholars, and faculty with opportunities to work on technical/research projects, an increase in the representation of these opportunities in scholarly literature would benefit funders, host institutions, and participants.^77^

Although previous publications about programs or outreach at national laboratories provide useful information about previous activities, many of these publications include data collected from students without documentation of informed consent or review/guidance from an Institutional Review Board (IRB); make claims about student learning that are not supported by the type of data collected or previous studies; or make sweeping generalizations about undergraduate education and/or student learning that may lead to inaccurate conclusions about program impacts.^78^ At the time of this writing we are not aware of any previously published studies that include data from program participants in DOE national laboratory programs aligned with IRB standards for human subjects research. This may be due to the fact that the IRB review has not taken place prior to collecting data from program participants or that this step was not documented as part of a manuscript.^79,80^ People have the right to be “respected, and to determine their involvement (or not) in research,” and studies that include student/participant data should include text to confirm compliance with institutional ethics guidelines.^80^ In studies about programs at DOE national laboratories, the lack of systematic investigations and/or commentary about human subjects protocols is problematic because it limits a) knowledge about how these programs compare to other similar programs, b) the extent to which scholars at other institution types can find and make contributions to what is already known about these programs, and c) productive collaborations with experts at DOE national laboratories on the subject.

### Theoretical Framework

Initially adapted from the social cognitive theory described by Bandura^81^, the **social cognitive career theory (SCCT) framework** was developed in order to understand the ways in which people develop their interest in a subject and make choices that ultimately impact their level of success in that field.^82,83^ The theory focuses on the relationship between certain cognitive-personal variables (**self-efficacy** and **outcome expectations**) with the supports and barriers an individual faces, and how this relationship influences the development of their career.^83^ In this study we applied SCCT to explore the impact of a particular **learning experience** on the academic and career trajectories of individuals who studied STEM while attending a U.S. community college.^17,84,85^ Valuable to our application of this framework, SCCT heavily weighs an individual’s belief in their ability to be successful in/on a given subject/task (**self-efficacy**) on their actual success in/on that subject/task. For example, if a student grows to believe that they are capable of being successful in STEM, this belief will work to shape their subsequent **interests** (which are fluid over time), **goals**, and **actions** related to STEM. In this way, the SCCT model proposes a connection between student experiences and perspectives with their future academic or career trajectory. **Self-efficacy** appears widely in the science education literature, related to numerous topics such as performance in STEM coursework, **career interest**, engagement in research, and success in graduate school.

Another component of the SCCT model relevant to this study are **outcome expectations**, which are an individual’s “beliefs about the consequences or outcomes of performing particular behaviors.”^86^ In contrast to **self-efficacy** (an individual’s perspective about their capabilities), **outcome expectations** involve predicting what will happen to them in the future, while striving to accomplish their **academic and career goals**. For example, a student may believe in their ability to learn and develop proficiency in biology, but they may not envision themselves being successfully admitted into a biology graduate program. In this example, a student may feel high **outcome expectations** related to pursuing a bachelor’s degree in biology, but low **outcome expectations** related to their application to graduate programs in this discipline. Studies suggest that academic and/or career-related **outcome expectations** may be a powerful influence over student behavior, even in the face of **contextual factors** that can be barriers to success, such as limited access to research opportunities and/or mentoring; experiences with discrimination and/or racism; or lack of career role models.^87,88^ This perspective is well-aligned with many studies that have used SCCT to understand the impact of science research experiences on persistence in STEM, an outcome which can be especially apparent for Black, Hispanic/Latinx, Native American, and female students.^89,90^

The pursuit of a STEM career requires “buy-in” from others, in the form of recommendation letters, information about funding, program, and employment opportunities, advisors or collaborators for new projects, etc. The SCCT model provides insight into the multiple ways in which students develop, sustain, and change their **career interests** in STEM, and are influenced by their experiences to make decisions to gain additional knowledge and professional experience over time.^82,83^ Students may have preconceived notions about working in their STEM field of interest from their interactions with others in school or the content of their coursework. However, these ideas may be misaligned with the actual experiences of people working in those fields. The SCCT model thus highlights the potential influence of regularly communicating with members of the STEM community on the process of clarifying and making decisions about possible career paths.^17,84^

Throughout the text, there are bolded terms that represent categories from the SCCT model (e.g., self-efficacy, learning experience, interest). We have highlighted these terms in bold, because they are the themes for which we have found connections between our data and the SCCT model. There are instances where these terms are referenced, but do not describe a connection supported by the data, and thus do not appear in bold.

### Role of SCCT model in STEM internships

We hypothesize that a STEM internship for community college students can serve as a **learning experience** aligned with the SCCT model, and elements such as interactions with mentors and role models can serve as **contextual factors**. Completing an internship could therefore influence the **interests**, **goals**, and **actions** of community college students, related to their academic and career trajectories. We recognize that a STEM internship is only *one* of many possible factors influencing career choice behavior (**actions**) for a particular student. However, previous evidence about the ways in which undergraduates are impacted by mentored professional development and research experiences suggest that **self-efficacy** and **outcome expectations** are impacted by participation in this type of **learning experience.**^17,28,35,91^ This is likely to be true especially when the nature of the work during the STEM internship relates to the academic courses taken at the community college and/or a students’ specific **academic and career goals**.

An internship hosted by an institution such as a DOE national laboratory gives students the opportunity to meet and interact with members of the STEM professional community *outside* of their college or university community. This increases the potential for social integration into the STEM community and increased awareness about the spectrum of career options in their field of interest. Thus, a STEM internship may serve as a mediating factor between community college students’ initial curiosity or **interest** in STEM and their graduation and **persistence** in the STEM workforce.

## Methods

### Internship Characteristics

Founded in 1999, the Community College Internship (CCI) seeks to encourage community college students to enter technical careers relevant to the DOE mission by providing paid technical training experiences at one of 16 participating DOE laboratories/facilities. This DOE Office of Science Workforce Development for Teachers and Scientists program is offered on a national scale, but this study focused on the CCI program at LBNL, managed by Workforce Development & Education (WD&E) at LBNL. Applications for CCI are solicited annually, and those selected are placed with a Mentor Group at LBNL – research mentors and colleagues – for whom they will work as “interns.” Ideally, in addition to teaching new skills to support intern development from novice to advanced scientists or engineers (e.g., read and understand primary literature, perform laboratory techniques), the Mentor Group engages in mentoring practices (e.g., career exploration), as well.^92,93^ During the program, interns spend the majority of their time each week working with their Mentor Group to learn new technical/research skills and apply these to a specific project. Interns also attend mandatory events hosted by WD&E. On the first day of the program, Orientation introduces interns to program elements and resources at LBNL. Intern Check-ins are small group meetings with the program coordinator to discuss intern experiences and address issues. Internship Meetings are opportunities for interns to interact with peers and guest speakers from various disciplines. Interns present their work to members of the LBNL community and other guests at a Poster Session. To the DOE Office of Science Workforce Development for Teachers and Scientists, interns submit written deliverables (e.g., paper, poster) and pre- and post-surveys online. Optional activities include tours; lectures and seminars; networking; and workshops. An exit survey is administered to all CCI interns by WD&E during the final week of their program, to allow program staff to make improvements.

Previous studies have shown that being paid through an educational program or research experience contributes to undergraduate academic success, self-esteem, self-efficacy, and feelings of being valued by the sponsoring group or organization.^94–97^ The financial compensation provided to CCI interns at LBNL in 2016 included a stipend of $800 per week, a housing supplement of $300 per week, and reimbursement for travel costs to and from LBNL. To be eligible for these financial benefits, CCI interns completed their “onboarding” forms, worked 40 hours per week (which includes project tasks with the Mentor Group, completing required training, and attending internship meetings) during the internship term, and submitted deliverables (e.g., surveys, technical paper).

### Study population

Our study population, referred to in this study as “CCI alumni,” consists of individuals who participated in (and successfully completed) the CCI program at LBNL between Summer 2009 and Fall 2016 terms. In total, 93 individuals completed the CCI program in this time frame, and data about demographic data collected is shown in Table 2. We collected survey responses from 43 CCI alumni, and conducted interviews with 12 of these individuals. At the time of their participation in CCI at LBNL, their primary academic majors were as follows: 15 (35%) in civil and/or mechanical engineering, 14 (33%) in chemistry, 5 (12%) in biology, 4 (9%) in physics and/or mathematics, 3 (7%) in environmental science, and 2 (5%) in computer science and engineering. Of the CCI alumni represented in this study, 40 (93%) attended one of the California Community Colleges, and the remaining 3 (7%) attended schools in Illinois, Massachusetts, and New York. More than half of the CCI alumni in this study attended one of the following schools, listed in descending order: Contra Costa College, City College of San Francisco, Diablo Valley College, College of Marin, Hartnell College, Ohlone College, and Sacramento City College.

**Table 2.**
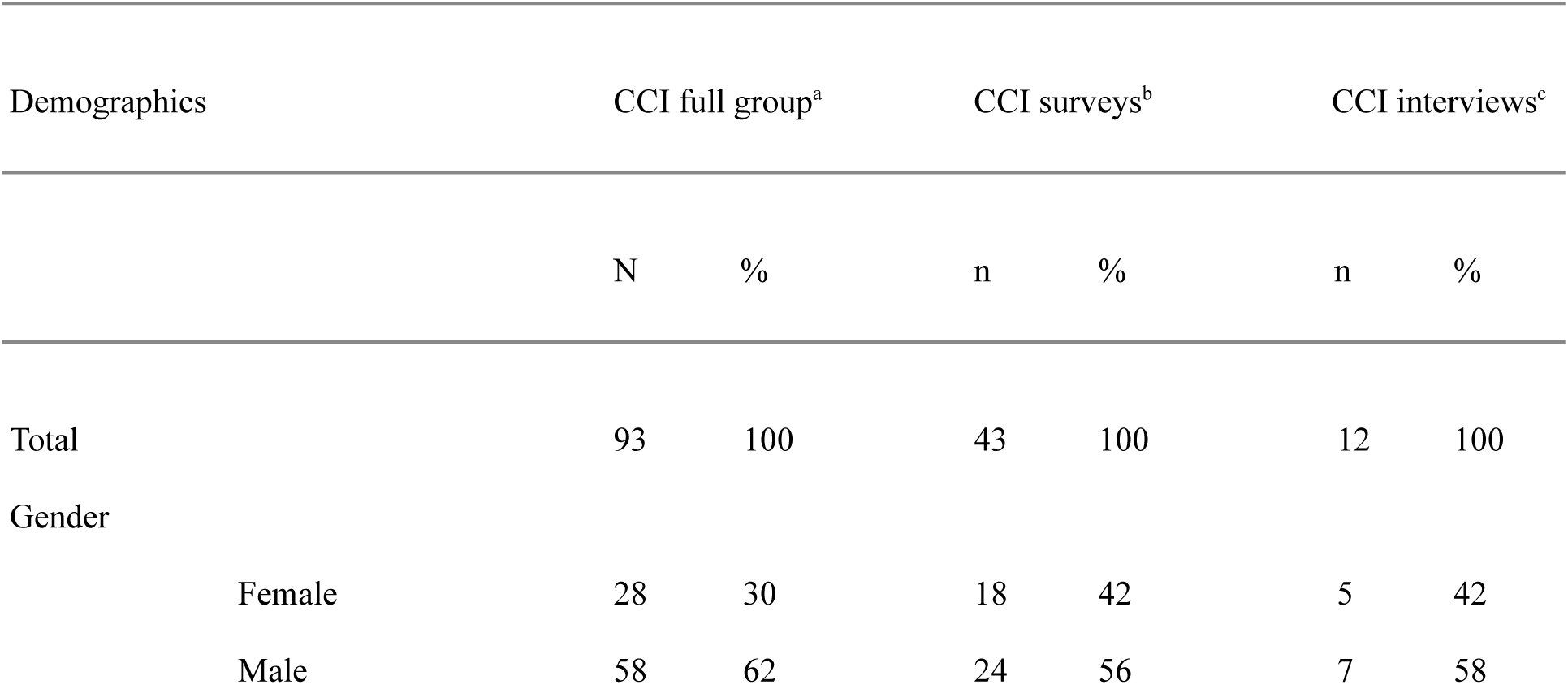
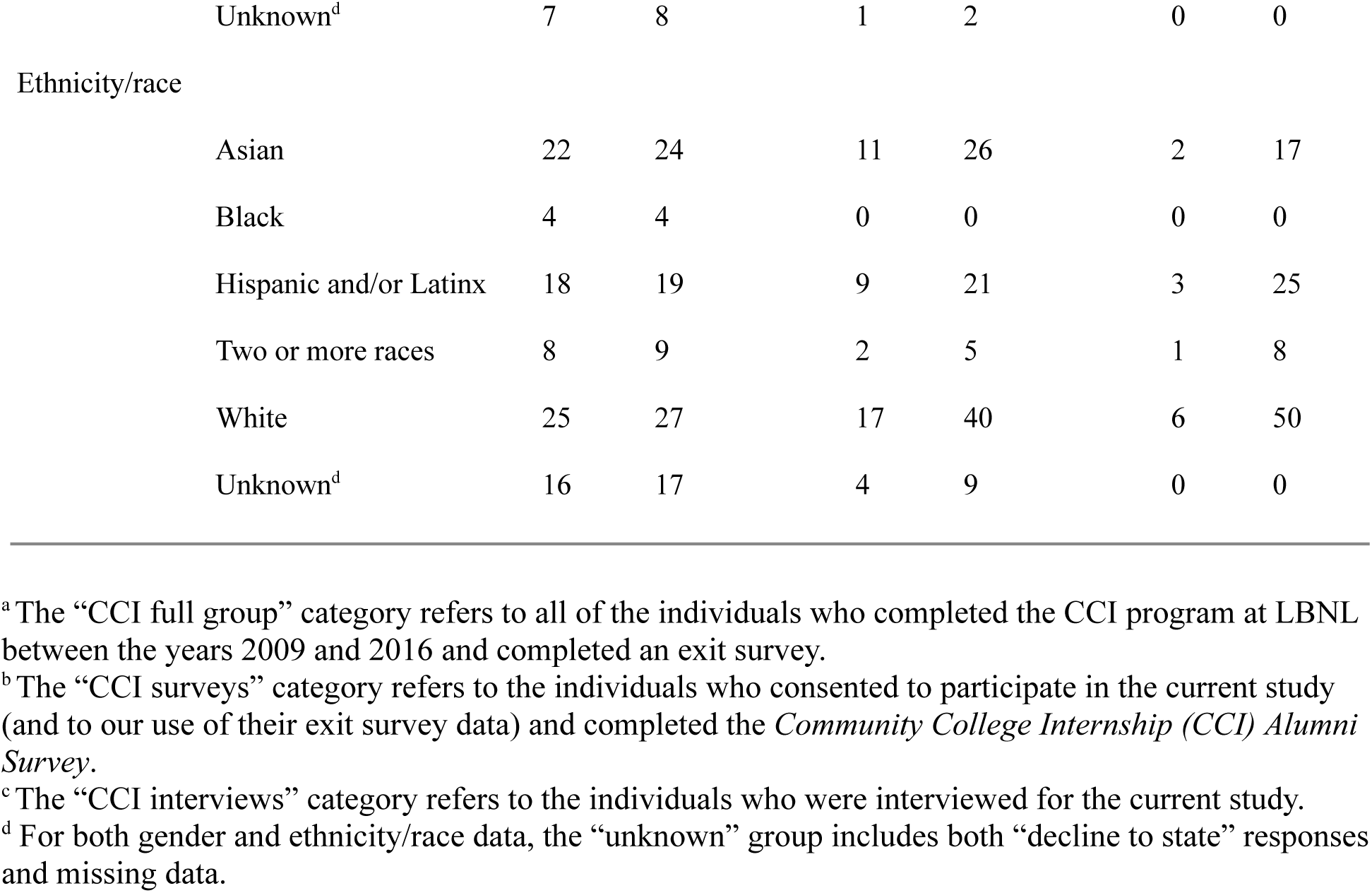
Demographic Information for CCI participants at LBNL.

We use the definitions of “science and engineering occupations” (e.g., life scientists, engineers, mathematicians) and “science and engineering-related occupations” (e.g., science teachers, laboratory technicians, laboratory managers) from the National Academies of Sciences, Engineering, and Medicine^98^ and National Science Board^99^ to define “STEM” in this context. Currently, 5 (12%) are graduate students, 36 (84%) have entered the STEM workforce, and 2 (5%) have entered the health workforce. Additionally, 20 (47%) have completed one or more A.A./A.S. degrees, 41 (95%) have completed one or more B.A./B.S. degrees, 11 (26%) have completed a master’s degree, 9 (21%) have – or have nearly – completed a Ph.D. in STEM, and 2 (5%) have completed a health-related degree (e.g., Ph.D., D.D.S., M.D., M.D.-Ph.D.).

### Positionality statement

The primary author (L.E.C.) is a woman who was born and raised in California, and grew up in a multicultural household in the U.S. with native-born and immigrant parents. Culturally she identifies as American and Middle Eastern. Her previous experiences as a community college student; participating in the CCI program at LBNL; conducting biological research as an undergraduate; and transferring to a California State University were helpful in establishing rapport with study participants. C.L.F. and L.E.C. have 14 and 13 years of experience, respectively, as practitioners working with a suite of internship programs at LBNL. A.M.B. has 28 years of experience working with and studying undergraduate research experiences for STEM majors, and L.E.C. has worked with A.M.B. on these projects for 8 years. Researchers E.W.L., J.J.S., and S.V.D. worked with WD&E internships as student assistants or employees. Researchers A.M., A.N.Z., and G.O.M. worked on this project as research assistants. Collectively, the authors of this work include individuals who identify as men, women, Asian, Black, Hispanic, Latinx, White, and mixed race. At the time of data collection and analysis, this team included undergraduates, post-baccalaureates, graduate students, faculty, and professionals in disciplines related to biology, chemistry, education, mathematics, medicine, physics, and public health.

As researchers our academic experiences, backgrounds, and identities prepared us to add context to the data we collected in this study and determine the best way to represent our findings. Community colleges have very little representation in the higher education literature, and authors of this study who attended community colleges worked to verify that this work did not perpetuate existing stereotypes about these institutions. Our previous work found that most studies about science research experiences do not a) report the proportion of participants from groups historically excluded from STEM fields, and/or b) present “disaggregated outcomes” for members of these groups.^56^ It became clear during the data collection process that our study participants’ experiences differed between groups, based on gender, race/ethnicity, and status as a first-generation college student. To ensure that our data analysis strategy would yield findings that highlighted these unique perspectives, co-authors who shared these identities were involved in exploratory conversations about the data we collected. This is discussed further in the “Data analysis” section. Described by Nakagawa and colleagues^100^, this study includes Method Reporting with Initials for Transparency (MeRIT).

### Data collection

An exit survey was administered to CCI participants during the final week of their internship (e.g., Summer 2009 interns completed exit survey in August 2009, Fall 2009 interns completed exit survey in December 2009), which gave us some baseline information about those elements of the program that were impactful to community college students at the time of their participation. This survey data was not available for students who completed CCI prior to 2009, so we chose to include study participants from 2009 and later.

Exit survey responses and published literature about community colleges, internships, and research experiences informed the development of the *Community College Internship (CCI) Alumni Survey* and interview protocol, which was written for use in the current study (S1 and S2 Figures). During recruitment in 2018, we found that some email addresses belonging to CCI alumni were missing or inactive, so the remaining CCI alumni were contacted through online platforms such as LinkedIn or Facebook. Consent forms and the *CCI Alumni Survey* were administered through Qualtrics. Semi-structured interviews were conducted a) in-person and recorded as audio files with a handheld recorder or b) using Zoom and recorded as both video and audio files. Interviews with participants were between 60 to 90 minutes in length. L.E.C. conducted the interviews, and field notes were taken by L.E.C. and S.V.D. The audio files were then transcribed and checked for accuracy by A.N.Z., E.W.L., G.O.M., J.J.S., L.E.C., and S.V.D.

The collection of *CCI Alumni Survey* data, interview data, and follow-up communication with study participants occurred between 2018 and 2021. This study was approved by the Institutional Review Board at LBNL (Protocol ID: Pro00023065) with the University of California, Berkeley, as the relying institution (Reliance Registry Study #2593). All contributing researchers have completed training in the responsible and ethical conduct of research involving human subjects, administered through LBNL or the University of California, Berkeley.

### Data analysis

When survey results were first obtained, A.N.Z. and G.O.M. organized open-ended responses into major categories, and these initial themes were used to guide discussions among our research team and with others who possess expertise regarding community college staff, faculty, and students in STEM departments (see “Credibility and trustworthiness” section). Data sets were organized by S.V.D. into individual folders for each study participant, to allow for review of all of the data collected from a particular individual for this study and guide our research team discussions. Survey responses and interview transcripts (referred to as “documents”) were analyzed using an approach that combined both grounded theory and content analysis: a) codes were generated based on the SCCT framework and literature related to professional development opportunities and research experiences for undergraduates; b) documents were read in full; c) sections aligned with the research questions were tagged; d) certain sections of each document were assigned one or more individual codes; e) codes that did not appear in our data were removed from the list of codes; and f) major themes identified in multiple documents were kept.^101,102^ After these major themes were identified, a first draft of the “Findings” section was written to summarize the ideas conveyed by study participants, supported by illustrative quotes. S6 Table 6 shows the list of categories identified from the SCCT model, the coding categories, and individual codes and sub-codes developed for use in this study. We discussed this draft to identify text that framed study participants’ experiences through a deficit lens, and made adjustments aligned with our asset-based perspective. For example, when study participants described their experiences and attitudes toward STEM before CCI, they often recounted stories in which they felt “clueless” about research and/or the work of scientists and engineers. We felt that it was important to balance the presentation of study participants’ stories “in their own words,” with the representation of their experiences in the context of the resources available to them, and the external factors impacting their perspectives. In the example above, we would include the word “clueless,” but worked to ensure that readers understood this to be the perception of study participants, and not our interpretation of their preparation or potential for success. Finally, we identified areas of the text which might be clarified or strengthened by conversations with others who have direct experience with the scenarios described by study participants. We then discussed these preliminary findings with community college students, faculty, and advisors and those who have served as technical/research mentors for community college students, which led to some additional insights. Finally, we discussed the content related to study participants’ unique experiences based on gender, race/ethnicity, and status as first-generation college students. Through these conversations, we identified the ways in which the data from these groups a) were characteristically different from the study population as a whole, b) could be supported by previously published educational research about these groups, c) could be framed through an asset-based lens, and d) furthered our understanding of the SCCT model applied to community college STEM majors. For example, the first draft of the current study addressed the importance of socializing during the learning experience, but did not include a connection between “warmth” in the social environment and the benefits of a learning experience as perceived by women. The identification of this connection led to some of the content that now appears as part of the “Gender” section (in Section 3). Coded documents were read again closely in order to modify/verify existing codes and apply new codes, and this led to multiple revisions of the “Findings” section text.

Scholars have called on educational scholars to report on the perspectives of Hispanic/Latinx community college students.^24,103,104^ When analyzing our data, we found that some study participants shared unique experiences connected to their **Hispanic and/or Latinx** identities. We acknowledge that the terms “Hispanic” and “Latino/a/x,” do not describe a singular racial, ethnic, or linguistic community.^105^ As shown in Table 2, approximately one-fifth of our study participants are Hispanic/Latinx. To protect the identities of our study participants, we have only used data in the “Findings” section that reference an individual’s self-identification as Hispanic or Latinx when that information was disclosed as part of an open-ended survey response or interview. Additionally, we have named the specific identity of one or more study participants when reporting a finding that connects to background, culture, or identity in our writing, instead of referring to a group as “students of color” or “underrepresented minorities.”^24^

### Credibility and trustworthiness

The topic of this study, CCI, is a national program funded by the DOE, and has multiple sites across the U.S. However, this study is focused on the CCI program hosted by one particular DOE national laboratory: LBNL. One benefit of examining the impact of the CCI program on participants who have all completed the program at LBNL is the reduction of institutional variability, as compared to a study comparing CCI across different sites.^106^

Used in several ways, triangulation was a key aspect of our approach to conducting this study.^107,108^ This study involves multiple data types, allowing us to employ data triangulation.^108,109^ Our primary method for collecting data was through the collection of survey responses. Conducting in-depth interviews allowed us to more clearly understand the survey responses by asking for clarification, collecting new stories, and allowing us to confirm or refine our initial interpretations of the survey data. We used analyst triangulation, in which multiple observers or analysts contribute their expertise to enhance the quality and credibility of our findings, in three ways.^108,109^ 1. There is a long history of assessment and evaluation of community colleges by people outside of the community college setting, which prevents those with expertise from contributing to knowledge about this system.^110,111^ As described in the “Data analysis” section, we consulted with current and previous community college students, community college and university faculty, university advisors who work with transfer students, and STEM professionals who serve as mentors. 2. During the data collection and analysis stages, multiple researchers read and discussed the data and the themes we constructed to present our findings. As we interpreted data from a particular study participant or group of study participants, authors of the current study with similar backgrounds, lived experiences, or identities were consulted. 3. Over the course of this study, we were in touch with CCI alumni at multiple time points for the purposes of recruitment into the study, scheduling interviews, and member checks to establish transparent relationships and rapport with this group.^112–115^ Although CCI alumni shared information about their lives that would eventually become data, we reduced the amount of “transactional” communication. Some CCI alumni reached out to check our progress or update our research team on their academic or career activities. They expressed gratitude for the opportunity to provide their perspectives, excitement for the publication of this work, and hope that it creates opportunities for community college students.

### Limitations

Selection bias is a common critique of studies focusing on the impacts of programs, internships, research/technical experiences, and other professional development opportunities. In this context, selection bias refers to the scenario in which a student makes the decision to apply to and/or has a greater than average chance of participating in a program as a result of their a) professional network, b) knowledge of its existence, or c) skill level in applying to the program (e.g., experience writing essays, relevant experience on their resume/CV, strong recommendation letters). We recognize selection bias as a potential limitation of this study, and thus cannot claim that participation in the CCI program directly caused the outcomes reported in this study. Instead, we make the case that participation in CCI influenced student perspectives about working in STEM fields, their own abilities and interest in STEM fields, and other related topics, aligned with the SCCT model. Our findings thus center around those outcomes that CCI alumni attribute to their participation in the program.

### Findings

All of the themes we constructed as part of our findings were informed by previous literature about SCCT, higher education, STEM persistence, and student learning during research/technical experiences and internships. Connections between our major findings and the SCCT model are summarized in Table 5. In Sections 1 and 2, our findings include major trends from study participants in the “CCI surveys” group (n=43; 42% female, 26% Asian, 21% Hispanic/Latinx, 5% two or more races, and 40% White) unless otherwise specified; see Table 2 for details. For example, a theme specific to Hispanic/Latinx study participants is described in the “Received support from and made connections with the LBNL community” section. Section 3 references a subset of the study population (n=16) that – through surveys or interviews – included stories from their lives to connect their experiences in STEM with their background, identity, etc. Although we gathered useful stories from surveys, the richness of our interview data (n=12) allowed us to develop common themes associated with gender, race/ethnicity, and “first-generation to college” status. The characteristics of interview subjects are shown in S1 Table.

## Section 1. Pre-program experiences, supports, and barriers

### Support from community college faculty, STEM groups, and peers

While attending community colleges, the faculty, staff, and student peers at their home institution were a key source of support for CCI alumni (Table 3). Aligned with recent studies about the supportive social environments at community colleges, 30/43 (70%) of CCI alumni described their positive experiences with small class sizes, community-building activities, and mentorship.^116^ Some compared the individualized attention they received at their community colleges with the inaccessibility of faculty at baccalaureate granting institutions after transferring. Although many alumni were originally not planning to apply to any STEM internship or other research opportunity, 21/43 (49%) of CCI alumni reported that they were supported to do so through a “nudge” from community college faculty members, STEM club leaders, program staff, or peers (Figure 1). We conceptualize the “nudge” as strong and deliberate encouragement to pursue a particular opportunity, despite student reservations and/or low self-confidence. They were first introduced to the idea of engaging in professional development opportunities during class, in conversations with peers, or while attending an event hosted by a STEM club. Later, the aforementioned individuals or groups would increase the intensity and complexity of support (e.g., reminders in class, announcements during club meetings, and writing recommendation letters).

> Something that helped a lot was the [STEM club] …That was key for me. Getting in a group with people that were open-minded and willing to connect and doing it together … You see them again, and again, and again in the same classes, and that was something that was really vital, having that support system there.

> … we had instructors that … had careers in industry or worked at these research centers, so they want to share what they learned … my instructor told me about CCI … and said, ‘You know, you’ve got a good shot of getting in. Would you be interested?’

**Figure 1.**
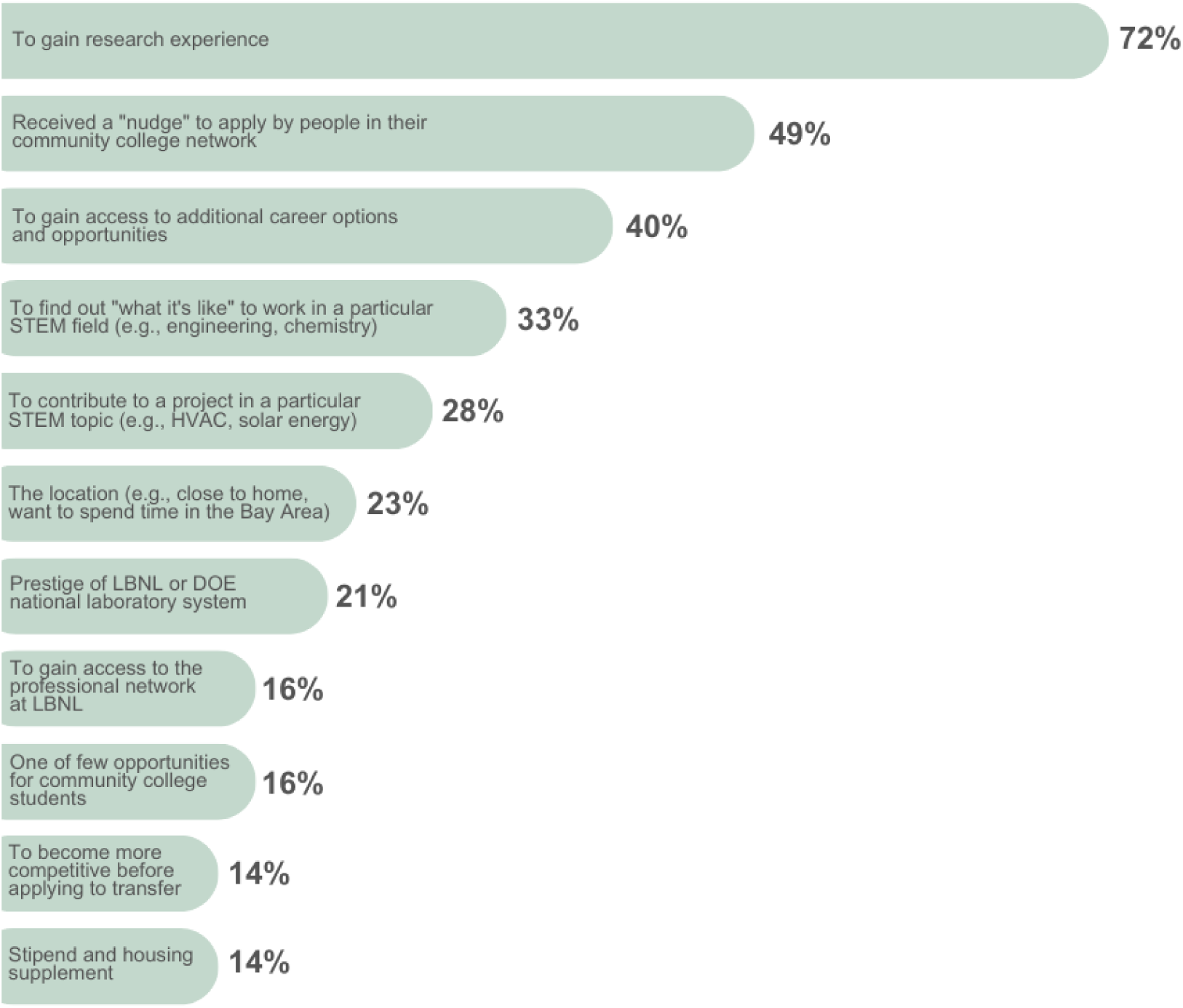
Primary reasons for applying to the CCI program at LBNL. These themes were generated from open-ended survey responses. Alumni applied to the CCI program between 2008 and 2015, and participated between 2009 and 2016. The percentage of CCI alumni (n=43) who listed each reason are shown. Respondents usually listed more than one reason.

**Table 3.** Experiences of CCI alumni as community college students taking STEM coursework before their participation in the program at LBNL.

| Theme | Sub-theme | Representative quote | % |
| --- | --- | --- | --- |
| Lack of knowledge about STEM research, jobs, and careers |  | “Before [CCI] I probably had no idea what a research question was ... on a zero to ten scale, before, it would be zero.” | 88 |
| Low expectations of success in a STEM career |  | “... my mentors told me to start applying to internships ... I had developed street smarts, so I had no issues in putting myself [into the] workforce, but when I exposed myself to science, I had uncertainties. I was worried if I’d find gainful employment that would make me happy.” | 86 |
| Support from community college faculty, STEM groups, and peers |  | “For me it was a really positive experience ... I had really good professors who, I think, kind of spent a little more one-on-one time with the students. ... I felt like I got to know my professors really well. It’s something I honestly would do all over again.” | 70 |
|  | Received a “nudge” from one or more people associated with the community college | “... the professors, the department, they mentioned it in classes, they mentioned this internship. They pushed [us] a little bit, but weren’t twisting everyone’s arm ... they made it really easy for [us] to get the information. Reminders about deadlines and stuff like that.” | 49 |
|  | Lack of support from family | “I am from a very traditional sort of community... going to college seemed like a joke [to them] ... without the [STEM club] at my community college, I would not be doing a STEM major. I would not have finished my degree.” | 21 |
| Few opportunities for community college students |  | “... people at my school, I have noticed that they don’t know about opportunities. When they find out I’ve had internships, they’re shocked! So, I think maybe other students are just going through this path, and they don’t know what else is out there ...” | 65 |
| Low expectations of being admitted into CCI |  | “I initially applied because I was encouraged by my community college mentors. I was extremely reluctant and felt that I did not compare well as a qualified candidate ... Never thinking that I would ever get accepted.” | 42 |

These themes are aligned with the social cognitive career theory (SCCT), and were developed from analysis of both open-ended survey responses and interview data. The proportion of CCI alumni (n=43) who reported experiences in each category are shown, with representative quotes for that category. Those study participants who completed a survey and an interview were counted only once. Many responses were multi-thematic.

For some CCI alumni, the lack of support from their families to pursue an undergraduate degree in STEM was a barrier to their academic success. 9/43 (21%) of CCI alumni described how this lack of familial support was challenging on an emotional level (e.g., self-doubt, embarrassment), or in terms of their understanding of what steps to take to pass their courses and/or transfer to a baccalaureate granting institution. Several Hispanic/Latinx alumni from rural agricultural communities explained that their upbringing did not prepare them to confidently select a major or obtain professional development opportunities, but that the social networks at their local community colleges were beneficial in helping them to achieve their goals. This is aligned with a study by Mireles-Rios and Garcia^117^, in which Latinx undergraduates described the importance of support from mentors and campus organizations to their academic success, confidence, and emotional well-being.

### Few opportunities for community college students

Alumni described a variety of goals for wanting to participate in the CCI program, including curiosity about research, a desire to apply the concepts learned in their STEM courses at the community college, or an interest to work on a particular STEM topic (e.g., clean energy, actinide chemistry, HVAC systems). However, 28/43 (65%) of CCI alumni articulated the lack of resources available to community college students. CCI was often described as being “one of the only” programs specifically for community college students, and one in which they were not in competition with students attending baccalaureate granting institutions as applicants.

> I’m glad that you’re doing research on this community college program. I think not enough opportunities are made available, or that we’re aware of [as students] … at [my school], it was unheard of. I know that <u>no one</u> prior to me had done the program.

### Lack of knowledge about STEM research, jobs, and careers

As community college students, 38/43 (88%) of CCI alumni, explained that they had little to no understanding of what it would be like to work in their STEM field of study before the program. Self-described as “clueless,” or having “no idea” about the work involved, many alumni shared stories about the moment when they found out that they had been accepted or in the weeks following. These were often connected to their fears, reservations, or predictions about the program. Others explained that although they had some conceptions of what scientists do (e.g., biologists use microscopes, physicists use complex equations), they did not understand the goals, processes, or rationale for this work. While attending community college, some individuals enhanced their interest in STEM through science-themed shows or magazine articles, but in retrospect, they explained that this media did not give them an accurate understanding of working in their STEM discipline.

> I was really surprised when I got it … I’m thinking, I have no idea what optics or x-ray beams are … I was talking to my chemistry professor, and I was like, “I don’t know what I just got myself into!”

### Low expectations of success in STEM

After taking some college-level courses in STEM subjects, but before completing the CCI program, 37/43 (86%) of alumni reported low expectations of their success in pursuing a STEM career. Their academic and career goals were vague at this stage. Some explained that they believed they could support themselves with a job, but felt unsure that they could be successful in the STEM workforce. For some, their own previously held misconceptions about community colleges led to lower confidence in applying to professional development opportunities. Although all of our study participants applied to and successfully completed the program, 18/43 (42%) of CCI alumni reported their initial low expectations of being admitted, and the belief that most professional development opportunities are meant for students attending baccalaureate granting institutions.

> … my mentors told me [about] programs that I should apply to, and felt it was out of my league. Students across the country from top schools were applying, so why should I apply?

> I don’t know, I feel like a lot of people think, “oh, <u>community</u> college,” and have a bad attitude. For me, looking back, [it] was an incredible decision for me, that completely changed my life.

## Section 2. Experiences during and after the CCI program

Shared through open-ended survey responses and interviews, CCI alumni explained what they gained from CCI (Table 4), and how the program shifted their perspectives about a number of topics. These gains were often contrasted with their perspectives as community college students before CCI, and stood out as being impactful to them years after completing the program.

> When I did my interviews for grad school, they asked me a question about internships. The first thing I talked about was my experience at LBNL, talking research projects, collaborating with professionals, problem solving on the spot. When I talk about it, my eyes light up! I’m in the moment, it really gets me excited. They see it, and they notice that, and they feel that, and then they know that I’m being honest and genuine. And they themselves understand that these experiences helped me get to where I am now.

**Table 4.**
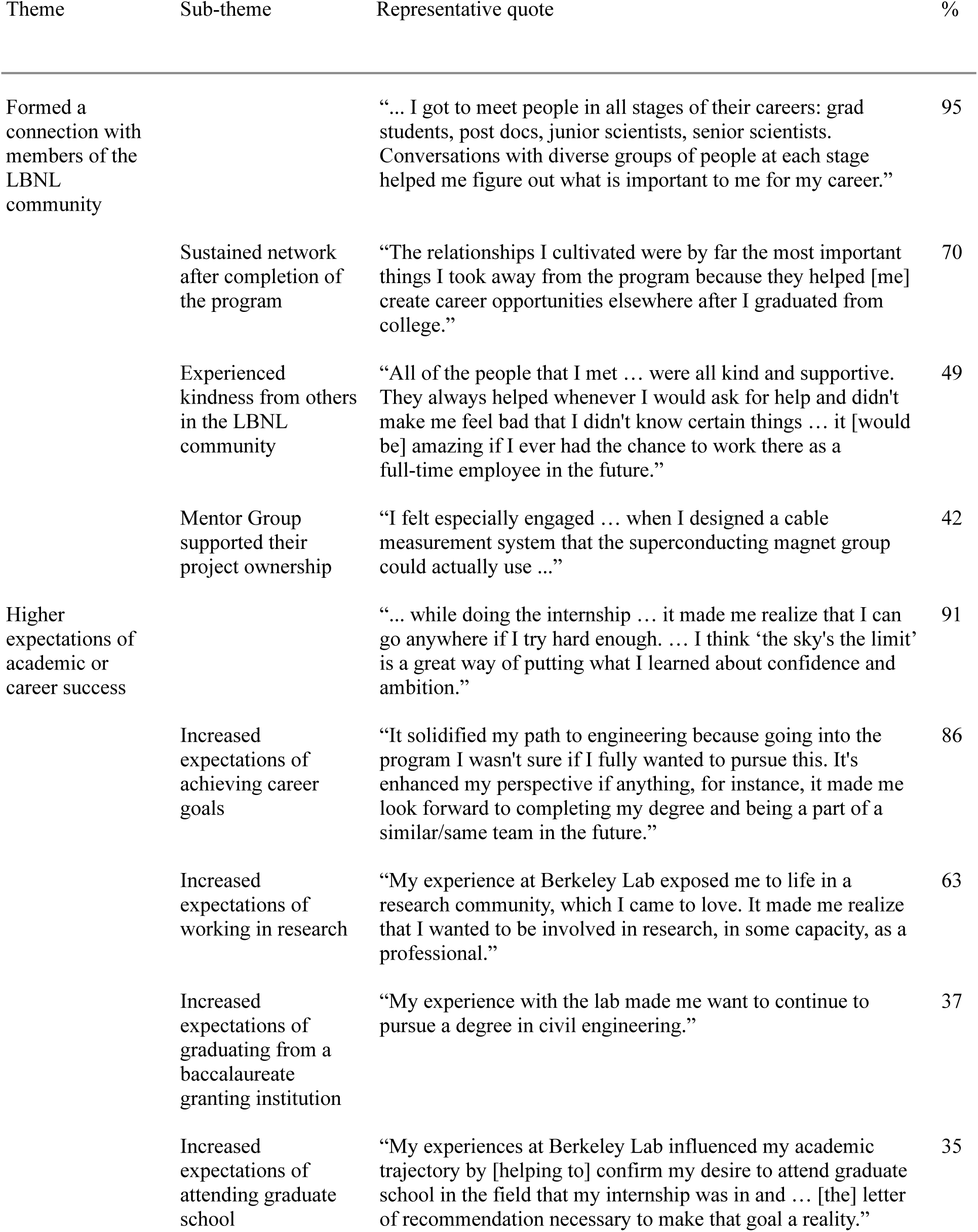
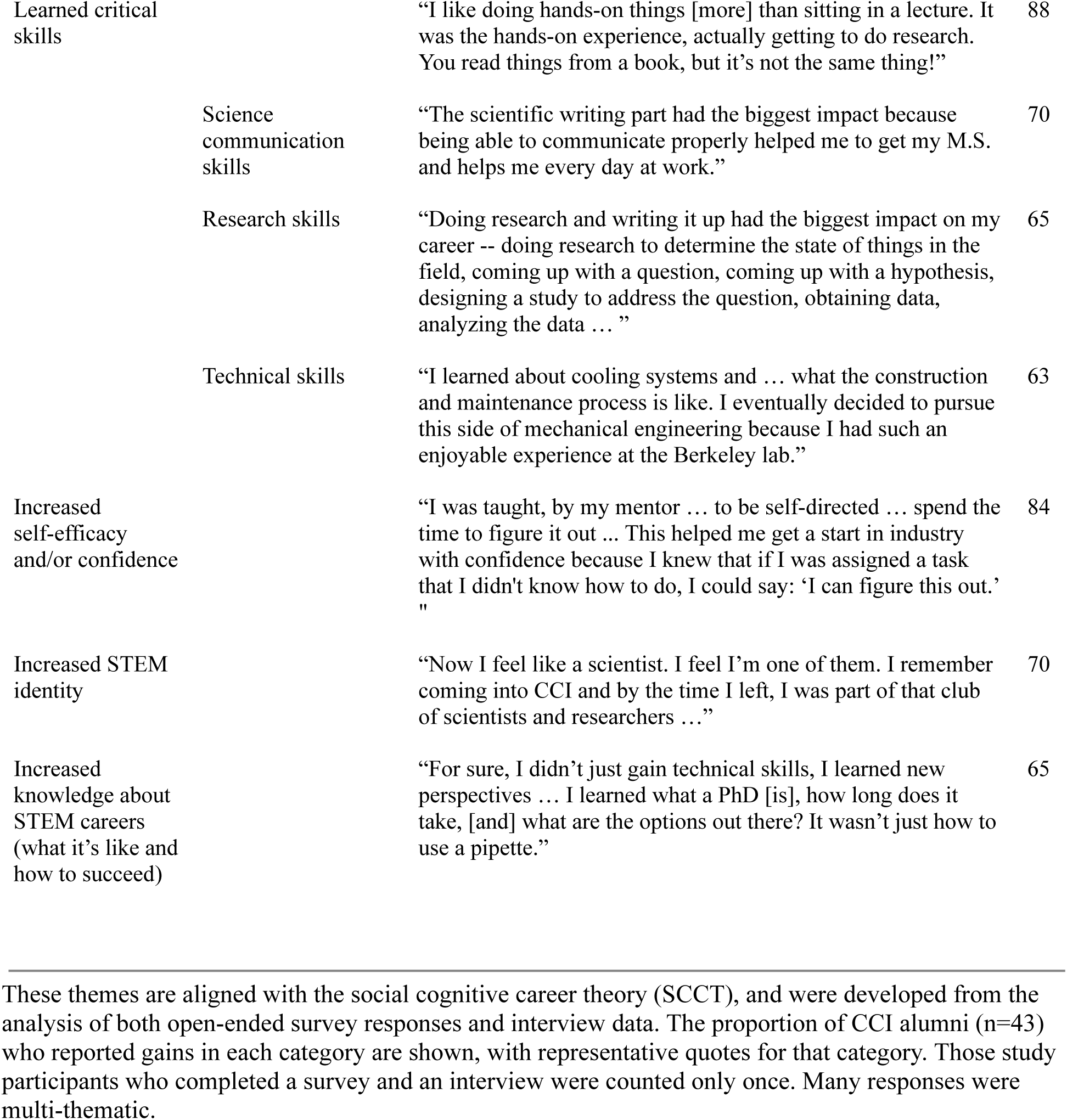
Most common benefits of participation in the CCI program at LBNL.

| Theme | Sub-theme | Representative quote | % |
| --- | --- | --- | --- |
| Formed a connection with members of the LBNL community |  | “... I got to meet people in all stages of their careers: grad students, post docs, junior scientists, senior scientists. Conversations with diverse groups of people at each stage helped me figure out what is important to me for my career.” | 95 |
|  | Sustained network after completion of the program | “The relationships I cultivated were by far the most important things I took away from the program because they helped [me] create career opportunities elsewhere after I graduated from college.” | 70 |
|  | Experienced kindness from others in the LBNL community | “All of the people that I met ... were all kind and supportive. They always helped whenever I would ask for help and didn't make me feel bad that I didn't know certain things ... it [would be] amazing if I ever had the chance to work there as a full-time employee in the future.” | 49 |
|  | Mentor Group supported their project ownership | “I felt especially engaged ... when I designed a cable measurement system that the superconducting magnet group could actually use ...” | 42 |
| Higher expectations of academic or career success |  | “... while doing the internship ... it made me realize that I can go anywhere if I try hard enough. ... I think ‘the sky's the limit’ is a great way of putting what I learned about confidence and ambition.” | 91 |
|  | Increased expectations of achieving career goals | “It solidified my path to engineering because going into the program I wasn't sure if I fully wanted to pursue this. It's enhanced my perspective if anything, for instance, it made me look forward to completing my degree and being a part of a similar/same team in the future.” | 86 |
|  | Increased expectations of working in research | “My experience at Berkeley Lab exposed me to life in a research community, which I came to love. It made me realize that I wanted to be involved in research, in some capacity, as a professional.” | 63 |
|  | Increased expectations of graduating from a baccalaureate granting institution | “My experience with the lab made me want to continue to pursue a degree in civil engineering.” | 37 |
|  | Increased expectations of attending graduate school | “My experiences at Berkeley Lab influenced my academic trajectory by [helping to] confirm my desire to attend graduate school in the field that my internship was in and ... [the] letter of recommendation necessary to make that goal a reality.” | 35 |
| Learned critical skills |  | “I like doing hands-on things [more] than sitting in a lecture. It was the hands-on experience, actually getting to do research. You read things from a book, but it’s not the same thing!” | 88 |
|  | Science communication skills | “The scientific writing part had the biggest impact because being able to communicate properly helped me to get my M.S. and helps me every day at work.” | 70 |
|  | Research skills | “Doing research and writing it up had the biggest impact on my career -- doing research to determine the state of things in the field, coming up with a question, coming up with a hypothesis, designing a study to address the question, obtaining data, analyzing the data ... ” | 65 |
|  | Technical skills | “I learned about cooling systems and ... what the construction and maintenance process is like. I eventually decided to pursue this side of mechanical engineering because I had such an enjoyable experience at the Berkeley lab.” | 63 |
| Increased self-efficacy and/or confidence |  | “I was taught, by my mentor ... to be self-directed ... spend the time to figure it out ... This helped me get a start in industry with confidence because I knew that if I was assigned a task that I didn't know how to do, I could say: ‘I can figure this out.’ ” | 84 |
| Increased STEM identity |  | “Now I feel like a scientist. I feel I’m one of them. I remember coming into CCI and by the time I left, I was part of that club of scientists and researchers ...” | 70 |
| Increased knowledge about STEM careers (what it’s like and how to succeed) |  | “For sure, I didn’t just gain technical skills, I learned new perspectives ... I learned what a PhD [is], how long does it take, [and] what are the options out there? It wasn’t just how to use a pipette.” | 65 |

These themes are aligned with the social cognitive career theory (SCCT), and were developed from the analysis of both open-ended survey responses and interview data. The proportion of CCI alumni (n=43) who reported gains in each category are shown, with representative quotes for that category. Those study participants who completed a survey and an interview were counted only once. Many responses were multi-thematic.

**Table 5.**
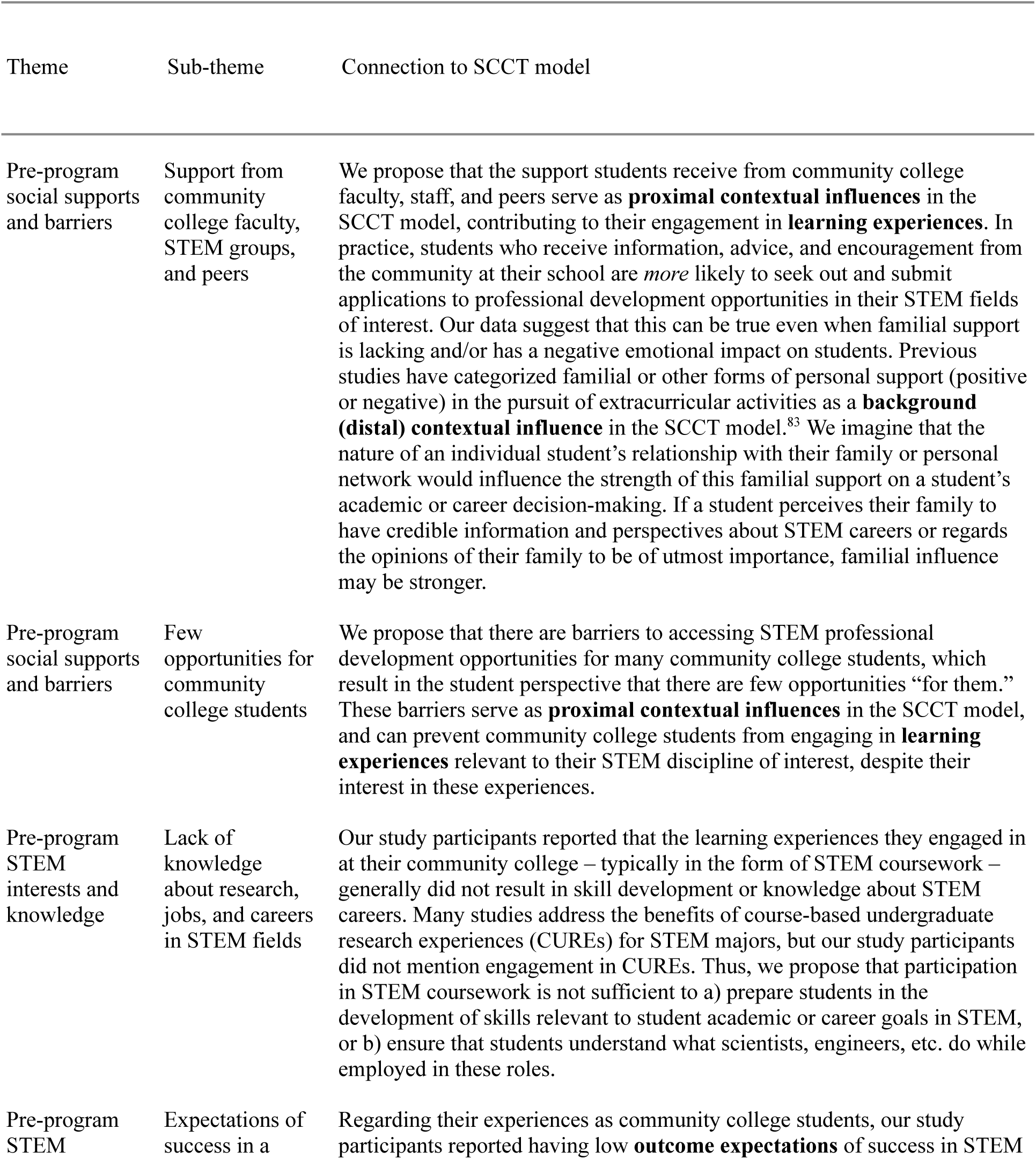
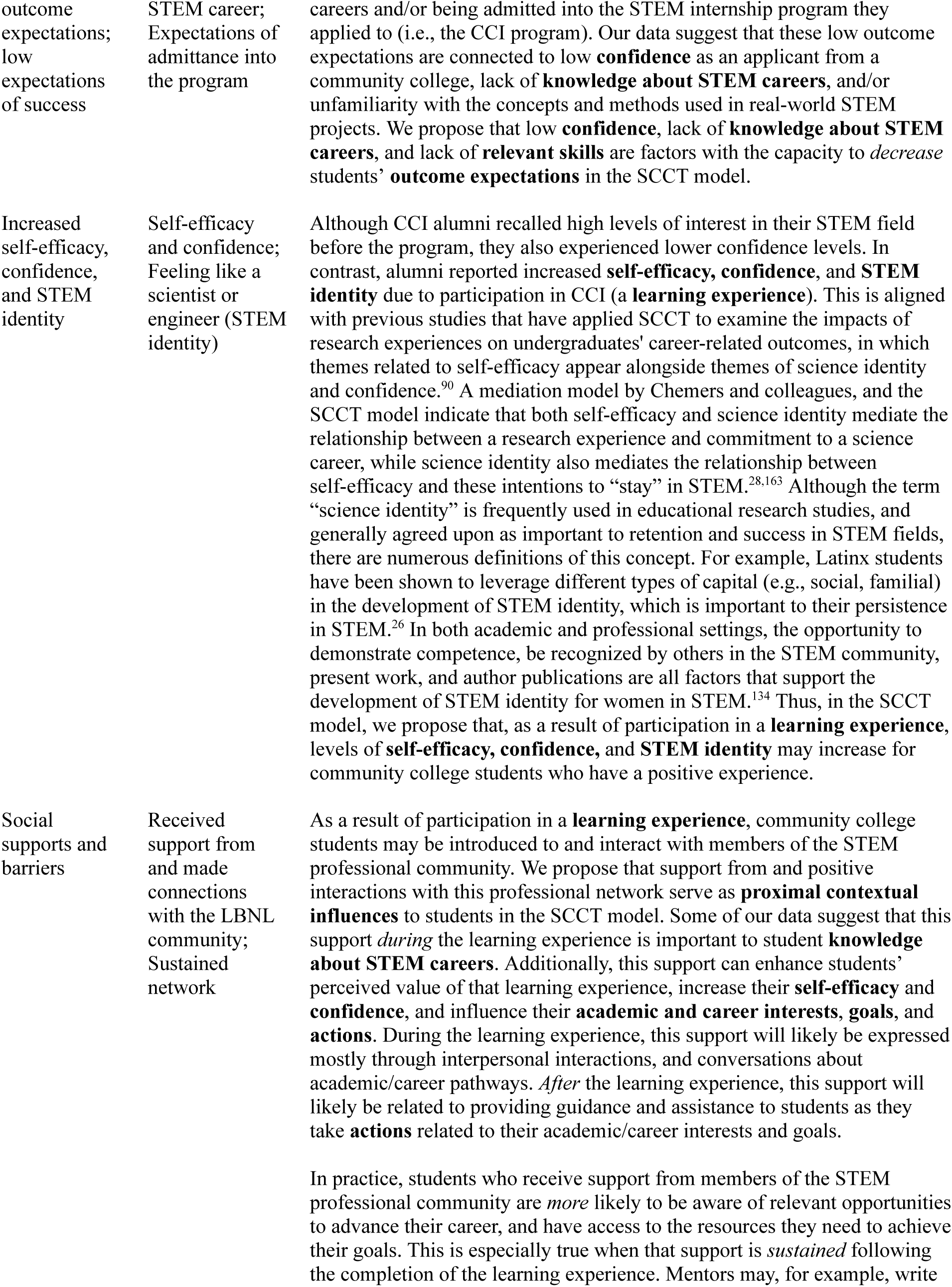
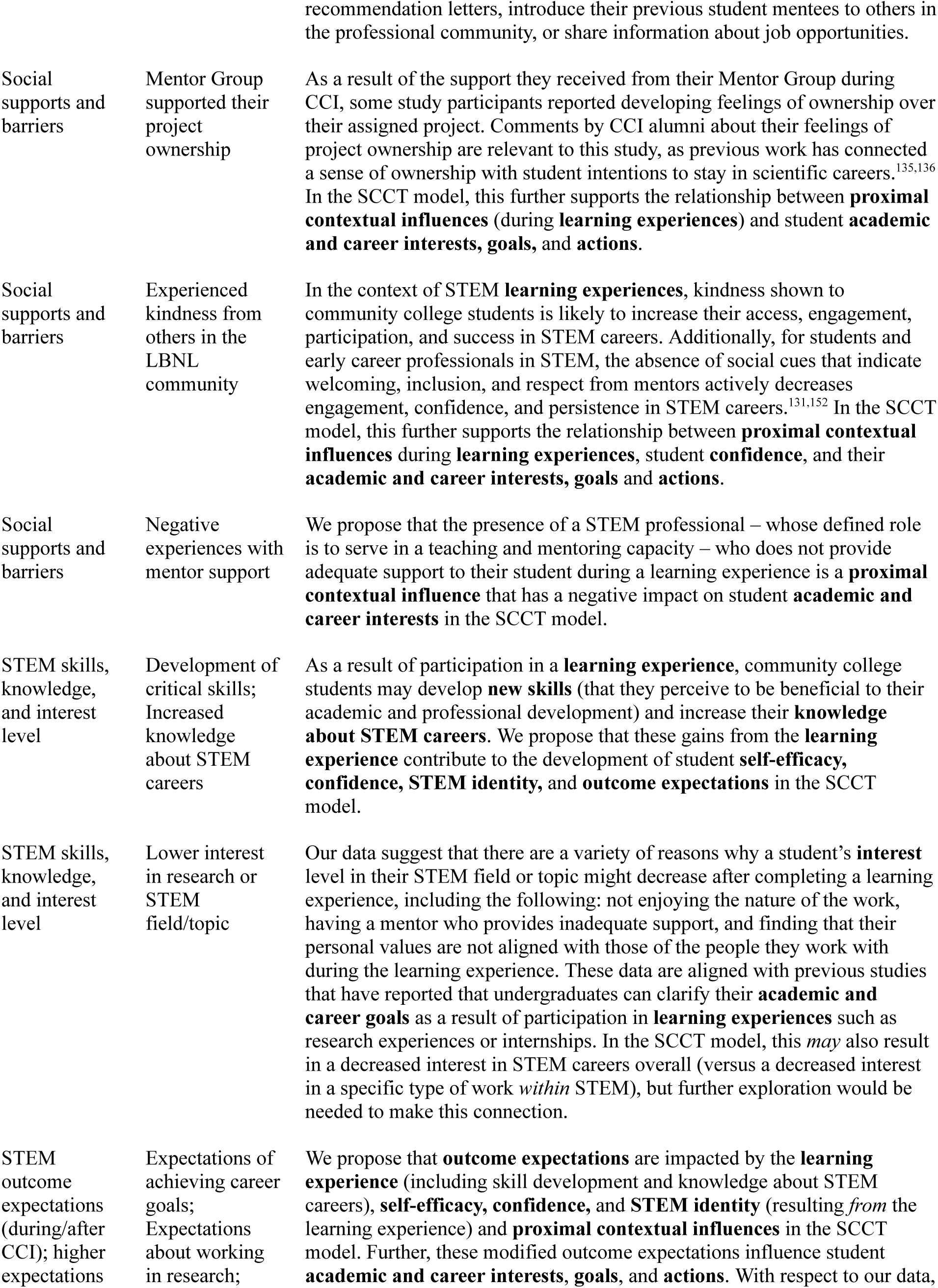
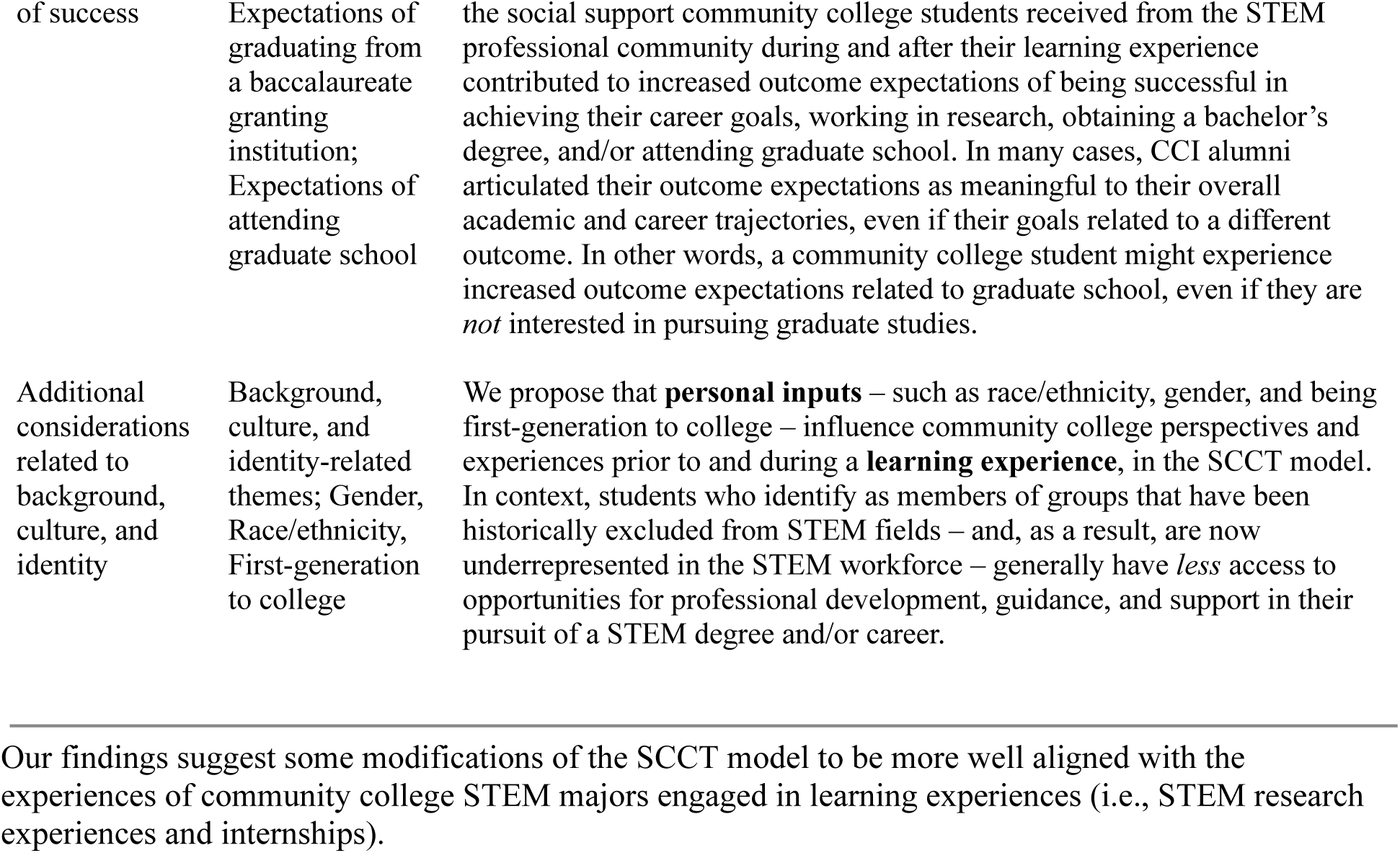
Connections between our findings and the Social Cognitive Career Theory (SCCT) model.

| Theme | Sub-theme | Connection to SCCT model |
| --- | --- | --- |
| Pre-program social supports and barriers | Support from community college faculty, STEM groups, and peers | We propose that the support students receive from community college faculty, staff, and peers serve as **proximal contextual influences** in the SCCT model, contributing to their engagement in **learning experiences**. In practice, students who receive information, advice, and encouragement from the community at their school are *more* likely to seek out and submit applications to professional development opportunities in their STEM fields of interest. Our data suggest that this can be true even when familial support is lacking and/or has a negative emotional impact on students. Previous studies have categorized familial or other forms of personal support (positive or negative) in the pursuit of extracurricular activities as a **background (distal) contextual influence** in the SCCT model.<sup>83</sup> We imagine that the nature of an individual student's relationship with their family or personal network would influence the strength of this familial support on a student's academic or career decision-making. If a student perceives their family to have credible information and perspectives about STEM careers or regards the opinions of their family to be of utmost importance, familial influence may be stronger. |
| Pre-program social supports and barriers | Few opportunities for community college students | We propose that there are barriers to accessing STEM professional development opportunities for many community college students, which result in the student perspective that there are few opportunities "for them." These barriers serve as **proximal contextual influences** in the SCCT model, and can prevent community college students from engaging in **learning experiences** relevant to their STEM discipline of interest, despite their interest in these experiences. |
| Pre-program STEM interests and knowledge | Lack of knowledge about research, jobs, and careers in STEM fields | Our study participants reported that the learning experiences they engaged in at their community college – typically in the form of STEM coursework – generally did not result in skill development or knowledge about STEM careers. Many studies address the benefits of course-based undergraduate research experiences (CUREs) for STEM majors, but our study participants did not mention engagement in CUREs. Thus, we propose that participation in STEM coursework is not sufficient to a) prepare students in the development of skills relevant to student academic or career goals in STEM, or b) ensure that students understand what scientists, engineers, etc. do while employed in these roles. |
| Pre-program STEM | Expectations of success in a | Regarding their experiences as community college students, our study participants reported having low **outcome expectations** of success in STEM |
| outcome expectations; low expectations of success | STEM career; Expectations of admittance into the program | careers and/or being admitted into the STEM internship program they applied to (i.e., the CCI program). Our data suggest that these low outcome expectations are connected to low **confidence** as an applicant from a community college, lack of **knowledge about STEM careers**, and/or unfamiliarity with the concepts and methods used in real-world STEM projects. We propose that low **confidence**, lack of **knowledge about STEM careers**, and lack of **relevant skills** are factors with the capacity to *decrease* students' **outcome expectations** in the SCCT model. |
| Increased self-efficacy, confidence, and STEM identity | Self-efficacy and confidence; Feeling like a scientist or engineer (STEM identity) | Although CCI alumni recalled high levels of interest in their STEM field before the program, they also experienced lower confidence levels. In contrast, alumni reported increased **self-efficacy, confidence, and STEM identity** due to participation in CCI (a **learning experience**). This is aligned with previous studies that have applied SCCT to examine the impacts of research experiences on undergraduates' career-related outcomes, in which themes related to self-efficacy appear alongside themes of science identity and confidence.<sup>90</sup> A mediation model by Chemers and colleagues, and the SCCT model indicate that both self-efficacy and science identity mediate the relationship between a research experience and commitment to a science career, while science identity also mediates the relationship between self-efficacy and these intentions to “stay” in STEM.<sup>28,163</sup> Although the term “science identity” is frequently used in educational research studies, and generally agreed upon as important to retention and success in STEM fields, there are numerous definitions of this concept. For example, Latinx students have been shown to leverage different types of capital (e.g., social, familial) in the development of STEM identity, which is important to their persistence in STEM.<sup>26</sup> In both academic and professional settings, the opportunity to demonstrate competence, be recognized by others in the STEM community, present work, and author publications are all factors that support the development of STEM identity for women in STEM.<sup>134</sup> Thus, in the SCCT model, we propose that, as a result of participation in a **learning experience**, levels of **self-efficacy, confidence, and STEM identity** may increase for community college students who have a positive experience. |
| Social supports and barriers | Received support from and made connections with the LBNL community; Sustained network | As a result of participation in a **learning experience**, community college students may be introduced to and interact with members of the STEM professional community. We propose that support from and positive interactions with this professional network serve as **proximal contextual influences** to students in the SCCT model. Some of our data suggest that this support *during* the learning experience is important to student **knowledge about STEM careers**. Additionally, this support can enhance students' perceived value of that learning experience, increase their **self-efficacy and confidence**, and influence their **academic and career interests, goals, and actions**. During the learning experience, this support will likely be expressed mostly through interpersonal interactions, and conversations about academic/career pathways. *After* the learning experience, this support will likely be related to providing guidance and assistance to students as they take **actions** related to their academic/career interests and goals.<br>In practice, students who receive support from members of the STEM professional community are *more* likely to be aware of relevant opportunities to advance their career, and have access to the resources they need to achieve their goals. This is especially true when that support is *sustained* following the completion of the learning experience. Mentors may, for example, write |
|  |  | recommendation letters, introduce their previous student mentees to others in the professional community, or share information about job opportunities. |
| Social supports and barriers | Mentor Group supported their project ownership | As a result of the support they received from their Mentor Group during CCI, some study participants reported developing feelings of ownership over their assigned project. Comments by CCI alumni about their feelings of project ownership are relevant to this study, as previous work has connected a sense of ownership with student intentions to stay in scientific careers.<sup>135,136</sup> In the SCCT model, this further supports the relationship between **proximal contextual influences** (during **learning experiences**) and student **academic and career interests, goals, and actions**. |
| Social supports and barriers | Experienced kindness from others in the LBNL community | In the context of STEM **learning experiences**, kindness shown to community college students is likely to increase their access, engagement, participation, and success in STEM careers. Additionally, for students and early career professionals in STEM, the absence of social cues that indicate welcoming, inclusion, and respect from mentors actively decreases engagement, confidence, and persistence in STEM careers.<sup>131,152</sup> In the SCCT model, this further supports the relationship between **proximal contextual influences** during **learning experiences**, student **confidence**, and their **academic and career interests, goals and actions**. |
| Social supports and barriers | Negative experiences with mentor support | We propose that the presence of a STEM professional – whose defined role is to serve in a teaching and mentoring capacity – who does not provide adequate support to their student during a learning experience is a **proximal contextual influence** that has a negative impact on student **academic and career interests** in the SCCT model. |
| STEM skills, knowledge, and interest level | Development of critical skills; Increased knowledge about STEM careers | As a result of participation in a **learning experience**, community college students may develop **new skills** (that they perceive to be beneficial to their academic and professional development) and increase their **knowledge about STEM careers**. We propose that these gains from the **learning experience** contribute to the development of student **self-efficacy, confidence, STEM identity, and outcome expectations** in the SCCT model. |
| STEM skills, knowledge, and interest level | Lower interest in research or STEM field/topic | Our data suggest that there are a variety of reasons why a student's **interest** level in their STEM field or topic might decrease after completing a learning experience, including the following: not enjoying the nature of the work, having a mentor who provides inadequate support, and finding that their personal values are not aligned with those of the people they work with during the learning experience. These data are aligned with previous studies that have reported that undergraduates can clarify their **academic and career goals** as a result of participation in **learning experiences** such as research experiences or internships. In the SCCT model, this *may* also result in a decreased interest in STEM careers overall (versus a decreased interest in a specific type of work *within* STEM), but further exploration would be needed to make this connection. |
| STEM outcome expectations (during/after CCI); higher expectations | Expectations of achieving career goals; Expectations about working in research; | We propose that **outcome expectations** are impacted by the **learning experience** (including skill development and knowledge about STEM careers), **self-efficacy, confidence, and STEM identity** (resulting *from* the learning experience) and **proximal contextual influences** in the SCCT model. Further, these modified outcome expectations influence student **academic and career interests, goals, and actions**. With respect to our data, |
| of success | Expectations of graduating from a baccalaureate granting institution; Expectations of attending graduate school | the social support community college students received from the STEM professional community during and after their learning experience contributed to increased outcome expectations of being successful in achieving their career goals, working in research, obtaining a bachelor's degree, and/or attending graduate school. In many cases, CCI alumni articulated their outcome expectations as meaningful to their overall academic and career trajectories, even if their goals related to a different outcome. In other words, a community college student might experience increased outcome expectations related to graduate school, even if they are *not* interested in pursuing graduate studies. |
| Additional considerations related to background, culture, and identity | Background, culture, and identity-related themes; Gender, Race/ethnicity, First-generation to college | We propose that **personal inputs** – such as race/ethnicity, gender, and being first-generation to college – influence community college perspectives and experiences prior to and during a **learning experience**, in the SCCT model. In context, students who identify as members of groups that have been historically excluded from STEM fields – and, as a result, are now underrepresented in the STEM workforce – generally have *less* access to opportunities for professional development, guidance, and support in their pursuit of a STEM degree and/or career. |

Our findings suggest some modifications of the SCCT model to be more well aligned with the experiences of community college STEM majors engaged in learning experiences (i.e., STEM research experiences and internships).

### Increased self-efficacy, confidence, and STEM identity

Before the program, including the period of time during which they were preparing their application, many CCI alumni explained that their background and/or lived experiences made it challenging for them to envision someone “like them” being successful at LBNL. However, 36/43 (84%) of CCI alumni reported that they experienced an increase in self-efficacy and/or confidence to work on technical projects and pursue their academic/career goals in STEM after the program. When CCI interns had opportunities to work in ways that pushed them beyond their comfort zone, the process of “getting through” these challenges was useful to their perception of their own capabilities.

> … [we] couldn’t figure out how to utilize [the] photomultiplier. We tried for over a day to figure out the proper orientation to apply voltage … It was incredibly frustrating and we were afraid [of looking] stupid … This moment was key… This experience … gave me confidence and ignited a passion for investigational projects.

Some CCI alumni recalled “always” having an interest in STEM, sharing stories from their childhood to illustrate the importance of these subjects in their lives. However, this early interest in STEM did not translate into confidence in their pursuit of a STEM career as community college students. Overall, 30/43 (70%) of CCI alumni reported “feeling like” a scientist or engineer during the program when they had the opportunity to spend time, collaborate, and complete projects with people employed in these positions. Some explained that it was meaningful to be paid a stipend by the CCI program, because this legitimized their roles as professionals in training.

> When I completed the flow schematics for an entire building, along with my team, and we had all the work in a giant portfolio. It really made us feel like we had completed our first engineering job.

> Yeah, just being in the cafeteria during lunch, that’s when I felt like a scientist. A strong emotional experience I [had] … taking the blue buses up to the hill … made me feel like a scientist every day of the summer program. It was just a phenomenal program that was a morale booster for me … I love that it felt like home.

### Received support from and made connections with the LBNL community

The most commonly reported gain, expressed by 41/43 (95%) of CCI alumni, was having formed connections with members of the LBNL community. These interpersonal interactions were often connected to a deepened understanding of STEM careers, related to the nature of a certain career pathway and/or the knowledge needed to take steps toward a particular career goal. Many CCI alumni shared their emotional response to working alongside people at LBNL who they perceived to be successful, values-driven, and accomplished. Several CCI alumni linked this professional network to their increased confidence in themselves as STEM majors upon returning to school after the program. One individual explained that having conversations with people at LBNL about physics – on the shuttle bus or in the cafeteria – gave them a “psychological boost” that was useful to them when they went back to school.

> It was very awe-inspiring … There’s all this peripheral learning that goes on. You interact with other scientists. I think it was really wonderful how the program organized all these events, opportunities to talk to different scientists to hear research talks from different scientists … just being in that environment was very beneficial.

For 30/43 (70%) of CCI alumni, this network was sustained after the completion of the program, leading to direct career benefits, including access to internships, jobs, graduate programs, and research collaborations. Examples of this continued support included receiving advice to inform career decisions; obtaining recommendation letters; new opportunities to collaborate with the team; being introduced to others; and learning about additional academic (e.g., programs, fellowships) or professional (e.g., jobs, internships) opportunities.

The kindness and socioemotional support they received from group members, program staff, and peers during the program was described by 21/43 (49%) of CCI alumni. In most cases, their experience of receiving kindness led to feelings of closeness, pride, engagement, and long-lasting positive feelings about the program and/or members of the LBNL community. Within the larger alumni group, 7/9 (78%) of Hispanic and/or Latinx CCI alumni shared examples of their experiences with kindness during the program. In addition to the aforementioned impacts expressed by other alumni, Hispanic/Latinx alumni shared how the receipt of kindness during CCI had a lasting impact on their impression of LBNL and their desire to work there again in the future.

> I worked with a wonderful mentor … [we] would talk about everything … marriage, to religion and metaphysical questions, we’d talk about science, basically anything you can imagine! … [My mentor] was truly a friend.

> … [my mentor] told me about the process, that scientists read through the application … [he] read through mine and said, “I have to meet her” … I still have that email saved.

During the CCI program, members of the Mentor Group engaged in teaching their interns new technical skills and how to approach problems in their field. Beyond this, 18/43 (42%) of CCI alumni explained that their Mentor Group trusted them to be responsible for some aspect of the CCI project after initial training was complete, and enabled them to take an active role in their own learning.

### Negative experiences with mentor support

In contrast to the sentiments expressed by most CCI alumni, 5/43 (11%) reported that they did not receive adequate support from their mentors during the program. In previous studies, some common characteristics of inadequate or negative mentorship include those mentors who are “too busy” to provide support, infrequently communicate, are overly critical, or show no interest in student technical or professional development.^118–120^ For three individuals in this study, their mentors practiced a “hands-off approach,” were not approachable, assigned their interns “menial” tasks, and/or seemed frustrated when interns needed assistance. The other two individuals explained that their mentors were not present or available to meet regularly during the internship. Even though their few conversations with their mentor were positive, their overall impression of their working relationship was negative. All five alumni in this group commented on their STEM interest; four were still interested in STEM after the program, but not in the field/topic they worked on during CCI, and the fifth individual was no longer interested in pursuing a career in STEM following the CCI program.

> I’m in that lab all day, and you know, [my mentor] is right there … she wasn’t necessarily the easiest person to walk up and tap on the shoulder that often … She kept it pretty formal … I didn’t want to bug her very much.

> I was a little disappointed in the work that was asked of us … they were utilizing their interns to do the work they did not want to do … we were only allowed to shadow [LBNL staff] … never work on a project with them or assist them in any way.

### STEM skills, knowledge, and interest level

As a result of participating in the program, 38/43 (88%) of CCI alumni reported that they learned skills valuable to their professional development. Specifically, 30/43 (70%) reported gains in scientific communication related to the preparation of written technical/research reports and public speaking experience; 28/43 (65%) reported gains in research skills, such as thinking like a scientist, organizing oneself for laboratory work, and analyzing data; and 27/43 (63%) reported gains in technical skills, such as producing drawings in AutoCAD, carrying out laboratory protocols, and learning a new coding language. Some individuals described how these new skills assisted them in their STEM coursework through a deeper understanding of STEM content knowledge or increased confidence when completing assigned projects. One individual learned finite element analysis and how to design a piece of equipment, both useful in their engineering studies and jobs after the program.

Most often through conversations with the Mentor Group or others in the LBNL community, 28/43 (65%) of CCI alumni expanded their knowledge about STEM careers. Some recalled instances where these professionals shared information about being successful and included personal details about their lives. These conversations seem to have been interpreted by alumni as microaffirmations – defined by Estrada and colleagues^121^ as subtle or ambiguous kindness cues – that they too could work at an institution like LBNL. Other CCI alumni were interested in learning about how to find a career path aligned with their personal goals and values, and found it meaningful to learn about the scientific and/or societal context of their project.

> It wasn’t just work, there was that actual mentorship. It wasn’t just like, “we’re splitting cells today,” or cleaning [something]. He was actually like, “this is my background, this is what I do. These are some things you could look into. Have you heard about this program?” … It wasn’t just work.

> STEM is hard. It’s like gymnastics or going to the gym … It takes dedication … mentors or professionals that basically look after women in STEM [are important] so that you don’t constantly feel like you’re struggling upstream alone.

After completing the program, 8/43 (19%) of CCI alumni reported that they were less interested in research, or the specific STEM field they were exposed to during CCI. One individual enjoyed working in STEM, but found a lack of alignment between their personal values and the values of the Mentor Group (and colleagues in the same department). They concluded that they were not interested in working in that field/topic, though they were still interested in joining the STEM workforce. Originally interested in graduate school, another individual became less interested in Ph.D. programs after the CCI program, because they learned that there are many technical roles available that require less training. Due to the “slow pace” of research, two individuals explained that the program allowed them to make an informed decision not to work in research; one currently works in a health field, and the other works at a DOE national laboratory in STEM in a technical role that does not involve research. A third individual explained that they were less interested in research after CCI, partly because their project was never well-described to them.

### STEM outcome expectations (during/after CCI); higher expectations of success

As compared with their experiences prior to participation in the program, 39/43 (91%) of CCI alumni reported an increase in their expectations of success in STEM academic programs and careers. Although alumni possessed a wide variety of career goals, 37/43 (86%) increased, broadened, and/or changed these goals due to participation in the CCI program. In some cases, this change related to the types of work they believed they were capable of doing, while others explained that their goals were “higher” than before. Some explained that they had identified jobs in the STEM workforce that they were interested in, but did not view themselves as potential candidates for these positions until after they participated in the program. Similarly, 27/43 (63%) of CCI alumni explained that the program helped them to maintain or increase their interest in a research-based career. Many CCI alumni explained this new interest in the context of “being exposed” to research, which involved learning about what research entails and gaining experience conducting research in an authentic setting.

> I am planning to add in a career step [and study] at the U.S. base in Antarctica. The CCI program really opened up my eyes to the possibilities that I have in which I never [knew] were so close.

As community college students, some (but not all) CCI alumni were certain of their desire to transfer to a baccalaureate granting institution prior to the CCI program. Related to this, 16/43 (37%) of CCI alumni linked their experiences in CCI with a new and/or strengthened desire to obtain a B.A./B.S. degree. Some individuals explained that their experiences made them more confident in transferring to particular universities or obtaining degrees in disciplines they had not previously considered. For 15/43 (35%) of CCI alumni, the idea of going to graduate school felt more attainable than it had before they participated in CCI, even for those who ultimately made the decision not to apply.

> Before I participated in the CCI program, I did not like molecular biology because [of] my previous professor … my PI and mentor were really helpful in explaining how the concepts of molecular biology were related to the research project. The opportunity changed my major decision.

> … before CCI, we were joking about how we were all going to get our Ph.D.s, but after, that was actually an option.

## Section 3. Additional considerations related to background, culture, and identity

In the field of undergraduate STEM education, many scholars have called for researchers to consider the ways in which students’ multiple identities can result in unique lived experiences, versus the examination of experiences based solely on a single identity.^122,123^ The concept of “intersectionality” was first introduced by Crenshaw as a response to the unique marginalization of Black women, as opposed to the experiences of Black people of any gender, or women of any racial/ethnic background.^124,125^ Studies exploring intersectionality in STEM have highlighted the value of storytelling in empowering students to share about the ways in which they believe their identities and lived experiences have impacted their STEM trajectory.^126–129^

To add context to their academic/career experiences, 16/43 (37%) of CCI alumni shared stories about how their upbringing, group membership, or culture impacted their experiences in STEM. The role of the mentor was a major theme present in both survey and interview data from alumni who self-identified as part of an “underrepresented” group. Despite being from a “different background,” mentors and staff who deliberately dedicated time to connect with interns about their personal lives were remembered as critical to interns’ positive experiences during CCI. This aligns with the concept of *personalismo* in Hispanic/Latinx culture, which describes how personal relationships are initially valued more than formal/institutional relationships, and critical to building trust (*confianza*) in an educational or professional setting.^117,130^

> Yeah, my mentor was a White man, but for me … he was truly a friend. [We] would talk about everything, basically anything you can imagine.

> [My mentor] would sit you down, and he would tell you anything you wanted to talk about. … I told him, I feel inadequate. I don’t know what I’m doing really, … And I remember he gave me a look, like he wasn’t prepared, because typically the interns that he gets, they already had an idea as to what they want to do. … he was a little surprised and interested … So, he started talking to me about his private life a little bit, sharing about some of the things he did … That was really helpful. … I know he had a lot on his plate, he always did. But, he kept his door open. He literally kept his door open.

### Gender

During interviews, 5/12 (42%) of CCI alumni described how their identity as a woman impacted their academic and professional experiences in STEM and expressed the importance of having access to a “warm” social environment in which they could interact with other women at LBNL during the CCI program. This included peers, mentors, and other staff.

> As a student I needed exposure to other STEM students who were equally as excited to do research, and CCI created that space during [group meetings]. Also, I needed [female] mentors since at my community college I was the only minority female in engineering.

Despite professional experience, entry into the STEM workforce, or completion of graduate training, some women shared their long-term struggle with identifying as scientists or engineers. One woman with a STEM graduate degree explained that she has never felt comfortable using these terms to describe herself. However, when asked to compare her role in the scientific community to people she would describe as “scientists,” she could not identify any differences between their professional activities and her own. She was comfortable with being called a “scientist” by non-scientists, but felt hesitant to use the term around others in the lab where she works.

> I still struggle with the word “scientist.” That’s not uncommon for women who are doing science. I would definitely say I am still learning. I’m a learner … I think the thing with [the term] “scientist” is that it feels like a bar so high, it’s something you’re always striving for, where you’re always being very careful about what you’re doing, and reading everything, and being very diligent about marking down what you’ve done …

### First-generation college students

Some of the alumni we interviewed described how being the first in their family to attend college (first-generation college students) impacted their experiences as undergraduates, and made it more difficult to access professional development opportunities. During interviews, 4/12 (33%) of CCI alumni shared stories in which they connected their status as being first-generation college students with recurring struggles to feel comfortable learning and working in STEM, even after earning undergraduate and graduate degrees in STEM. This group of alumni are diverse in gender, race/ethnicity, and STEM field of interest. They all made references to the fact that they did not receive advice and/or support from their personal social network when making academic- and career-related decisions.

> Being first-generation … there’s no one really before me that can give me tips on how to navigate this world … At LBNL, I was feeling a little inadequate … Like, “I shouldn’t be in a place like this. There’s Nobel Prize winners around! We’re developing special bacteria that can eat through plastics, detecting neutrinos, and all this fancy incredible stuff.” … And I’m like, “I’m from a farming community.” And I felt that. I told [my mentor], “I feel inadequate.”

### Race and ethnicity

During interviews, 4/12 (33%) of the CCI alumni shared stories about how their racial or ethnic identities impacted their interpersonal relations with educators and STEM professionals. Some CCI alumni explained that the “diversity problem” in STEM made it challenging for them to envision being successful in STEM long-term. Several Hispanic/Latinx alumni explained how being “the only one” like them and facing racial discrimination led to lower confidence during school and when considering professional development opportunities.

> You go into a research facility and you see that there’s not too many people that speak like you, … they’re not Latino … when I went back to school, [I was dealing with] those negative thoughts.

One individual, who is Latino, described their early interest in working as an engineer. However, “those dreams dissipated” when they were repeatedly dismissed by most of their K-12 teachers, and became accustomed to frustration and disappointment in the classroom. Another individual shared how her early educational experiences inspired her to serve as a role model for others who may have experienced discrimination and bias in class and from society. Below she describes microaggressions – “small acts of aggression” that can cause self-doubt and psychological harm to their recipients – which can make students feel as though they have “prove” that they belong in STEM.^131,132^

> I gotta identify myself as Latina … cultural identity is a part of me, full-time. As a Latina, it’s important for me to represent. There’s not a lot of people like me in this school … We feel a lot more pressure … I remember as a young high school student, comments from teachers saying that they were surprised I was doing so well. “You’re a smart Mexican!” You hear that kind of crap … I don’t want to make it sound like I’m doing something important, because I don’t feel like I’m trying to be special … But, I am cognizant of the fact that, if I fail, or if I do poorly, I’m making it harder for people like me.

## Discussion

Although there is a great deal of literature linking technical/research experiences to persistence in STEM fields, very few studies examine participant perspectives and/or outcomes beyond the first few years after such an experience.^13,14^ In this study we connected the experiences of community college students – before, during, and after a STEM internship – with their academic and career activities in the years following the internship.

### Internships at DOE national labs integrate students into the STEM community

Due to the striking underrepresentation of DOE national laboratories and community colleges in the higher education literature, we used this study as a way to investigate both of these topics. Learning, collaborating, and spending time with STEM professionals outside of their school was impactful to community college students who previously struggled to imagine what scientists and engineers “do” at work. Many of the CCI alumni in this study recalled aspects of their experiences that are unique to working at LBNL — such as spending time in the cafeteria or riding the shuttle — that became important to them over the course of their internship. As these activities became familiar to the interns, so too did the idea that they were a part of the institutional community. Many CCI alumni reflected on the unique opportunity to explore, learn, and work at a DOE national laboratory during their internship. Some found it valuable to collaborate with experts in their field while others reflected on the benefits of having access to specialized research centers and powerful technology.

Scholarship about “ownership of learning” suggests that certain learning environments can create or strengthen excitement and motivation about the topic of study.^133^ In the current study, most CCI alumni attributed an increase in their STEM identity to the novel experiences of being involved in the activities of a scientist or engineer, including opportunities to apply their newly-learned skills, present their work, and be recognized by others as colleagues. Some found it valuable to connect their CCI projects to societal or scientific impacts, especially if they pursued a STEM career to make a positive impact on others. Others felt deep connections and commitment to their CCI projects when their Mentor Group gave them responsibility and ownership over some aspect of the work. This aligns with previous work about how opportunities to showcase one’s competence as a scientist through research progress and interactions with others can increase self-recognition as a scientist^14,134–136^; how STEM projects framed as beneficial to society can support STEM learning and retention for students with communal goals^93,137^; and the value to students in developing project ownership, a commitment and personal connection with a project.^133^ Our findings suggest that completing a STEM internship at a DOE national laboratory can produce outcomes that are comparable to other STEM research experiences or internships for undergraduates.

### Many students are looking for opportunities close to home

Many community colleges are disconnected from the STEM professional community and promote employment over professional development opportunities that would support the development of STEM identity in students.^50,138,139^ At the time of their participation in CCI, more than 90% of our study population were residents of the same state where LBNL is located (California), and most were attending community colleges located within 100 miles of the LBNL Main Site in Berkeley. This aligns with previous findings about community college student preference for completing professional development opportunities located within a “comfortable distance.”^140–143^

### Harmful stereotypes decrease student confidence

Our study participants identified stereotypes about community college students, from the media or harmful “comments” made by others. The academic community perpetuates negative stereotypes about community college students, which can have harmful impacts on community college student retention and belonging in STEM fields.^64^ Similarly, television shows, films, and books produced in the U.S. rarely include depictions of community college students and/or inaccurately portray them as “mediocre” and “unmotivated.”^144–146^ Although our study participants did not agree with these negative depictions, these biases negatively impacted their confidence to apply to internships or other professional development experiences outside of school. Our findings align with previous studies about the impact of negative narratives about community colleges on students’ “thoughts, beliefs, values, and behaviors,” even when they have positive perceptions about community colleges themselves.^147,148^

### Application selection processes can be updated to increase access

Our results suggest that community college students feel that most opportunities for professional development are not “for them,” especially when competing with students from more schools that provide more support with finding opportunities, writing personal statements, and obtaining recommendation letters. This aligns with previous work that has highlighted the unique challenges community colleges face with supporting their students to engage in STEM research, who are often unaware of research opportunities, believe that they are not qualified to apply, and are faced with implicit bias when they do apply for internships and research programs.^138,149^ Although many technical or research opportunities are offered to provide undergraduates with the chance to *gain* experience, studies suggest that “cultural biases of academic research” lead many STEM professionals to select those applicants who already *have* relevant experience or extracurricular activities for these positions.^150,151^

### Kindness supports inclusion and persistence in STEM

For students and early career professionals, positive and supportive mentor-mentee relationships in a particular professional environment contribute to an increased desire to remain in a similar career pathway.^152^ Multiple scholars argue that Black and Hispanic/Latinx students and professionals do not receive adequate support needed to obtain their academic/career goals, and call for additional studies on the subject.^153–157^ Throughout this study, we have reported examples of practices that led to community college students feeling as though they were capable, competent, and prepared to pursue a STEM degree or career. Previous studies about students from groups historically excluded from STEM have reported that a) mentoring relationships are necessary to retain these students in STEM careers, b) these students receive less mentoring overall, c) mentors with a similar background can be effective, but they are often over-burdened, and d) well-intentioned mentors can inadvertently harm these students through practices (e.g., biased selection of applicants, colorblind mentoring) that reproduce inequities faced by these groups in the past.^34,157,158^

Although kindness is valued by all types of students, studies have shown that kindness cues in the form of macro- and microaffirmations can contribute to feelings of social inclusion, and persistence in STEM for Black, Hispanic/Latinx, Native American, and low-income students.^121,159^ Two related studies that included more than 2,200 undergraduates majoring in life sciences attending baccalaureate granting institutions found that negative social interactions, feeling excluded or unwelcome, and witnessing unfair treatment – such as favoritism – were factors that led to students wanting to *leave* their group.^118,160^ For these students, a positive environment and experiencing kindness led to the opposite result, and they were more likely to *stay* with their group. Similarly, a study about female undergraduates majoring in engineering found that microaggressions led to feelings of exclusion, frustration, and a desire to limit social interactions; examples included people showing surprise that a woman would study engineering; having to prove to others that they are qualified to be in engineering learning environments; tokenization as one of few women in engineering; and overhearing inappropriate jokes told by colleagues/peers.^131^

### Advancing the SCCT framework

CCI alumni reported that **learning experiences** (including STEM coursework) they engaged with at their community college did not result in **skill development** or **knowledge about STEM careers**, and they had low confidence and outcome expectations. Their stories indicate that perceived barriers (e.g., biases against community college students, few opportunities) act as proximal contextual influences that reduce the likelihood of applying to a learning experience, but support from faculty, STEM clubs, and peers are proximal contextual influences that increase that likelihood. Many CCI alumni received a “nudge” to apply to the program from community college faculty, STEM club leaders, program staff, or peers, which was especially critical for those who experienced discrimination in K-12 educational settings or lacked familial support for their academic goals. A recent study identified types of individuals who influenced community college women of color during their pursuit of careers in STEM; family members, college faculty/staff, and K-12 educators were most commonly named as positive influences. ^161^ Notably, two of their study participants named K-12 educators as negative influences, who “tried to block opportunities for them to advance their education,” which aligns with our findings.^161^ The original SCCT model and subsequent iterations did not include a link between **proximal contextual influences** and **learning experiences**, but our data connect these two concepts (Figure 2). In other words, many of the students who participated in the CCI program would have been less likely to apply to the program without the support from their community college network.

**Figure 2.**
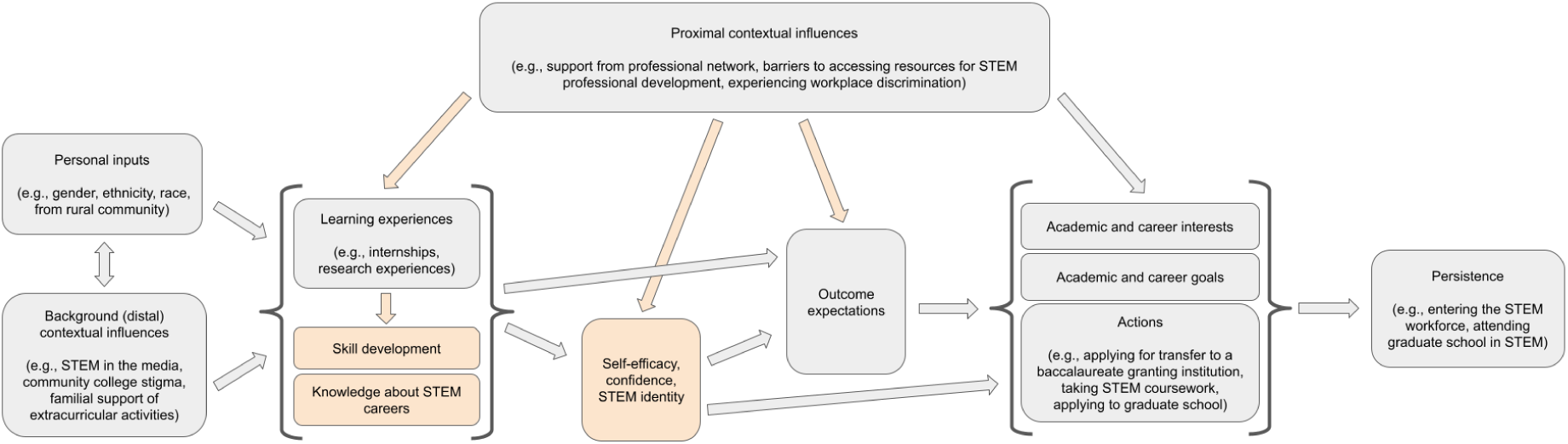
Model of how Social Cognitive Career Theory (SCCT) influences community college student participation in learning experiences and subsequent persistence in STEM fields. This was developed for the current study based on the original model of how basic career interests develop over time^82^ and later iterations applied to STEM learning experiences for undergraduates.^17,85^ Boxes highlighted in orange were added to previous models, based on data from Community College Internship (CCI) alumni in the current study.

Recent studies about the SCCT model suggest an indirect effect of **proximal contextual influences** on goals related to STEM careers through **self-efficacy** and **outcome expectations**, and other potential modifications to the model in its original form.^91,162^ Nearly all of the CCI alumni surveyed reported the two largest gains from participating in a STEM internship as a community college student to be a) the connections they formed with members of the STEM professional community (**proximal contextual influences**) they interacted with during the internship, and b) increased expectations that they would be successful in their academic or professional pursuits (**outcome expectations**). This professional community allowed students to learn more about STEM careers during CCI; supporting them in achieving their goals; and provided kindness – all of which resulted in increased student self-efficacy and confidence. These findings allow us to extend the SCCT model by including a direct link between **proximal contextual influences** and **self-efficacy** and/or **confidence**.

Mentors, colleagues, peers, and program staff continued to take actions to provide support in the form of advice, recommendations, additional internships and jobs, future collaborations, publishing papers together, etc. As described in the “Study Population” section, of the 38 CCI alumni in this study who have entered the workforce, 36 (95%) are working in STEM fields. Further, those who stayed in contact with the LBNL professional community after the CCI program were regularly reminded of their ability to be successful. These **actions** were thus connected to STEM **persistence**. Additionally, alumni reported that learning new research and/or technical skills and developing proficiency in science communication during CCI were beneficial to their academic and career trajectories and feeling like a scientist or engineer. Previously, the SCCT model did not include **skill development** or **knowledge about STEM careers**, although these are well-known outcomes of professional development opportunities for students. They have been added to the SCCT model in this study, because our data suggest that these are critical to increased **self-efficacy, confidence, identity,** and **outcome expectations**. Although we understand them to be individual concepts, there are a large number of studies that link **self-efficacy, confidence,** and **STEM identity** together, and our data suggest that all three of these factors are closely related.^28,30,35,54,90,163^ Thus, we have grouped these three concepts together in our proposed updates to the SCCT model, as applied to community college STEM majors who completed a STEM internship.

Some alumni expressed frustration when they felt like “the only” person from a particular background/identity in a particular group/setting, especially when this situation repeated itself over time. However, those mentors from a different race/ethnicity than their mentees who created space to have conversations about personal topics created feelings of trust and closeness. Aligned with the concept of “authentic care,” our findings indicate that mentors who were perceived to prioritize the care of mentees over project-related outcomes (e.g., finishing tasks, generating data) made a deep and long-lasting impact on their mentees.^164^ Women who participated in the CCI program reported the benefits of experiencing a “warm” social environment and opportunities to interact with other women. Some Hispanic/Latinx alumni shared stories about the discrimination they experienced from K-12 educators and schools, which negatively influenced their **confidence** to apply to the CCI program and their expectations of success in a STEM career. Combined with knowledge from previous studies about SCCT^17,85^, our data suggest that **personal inputs** (e.g., race/ethnicity, socio-economic status; first-generation to college) play a role in the pursuit of a STEM degree or career, and in their overall perspectives before, during, and after a **learning experience**.

Community college students may be actively exploring and negotiating their relationship to STEM as a career pathway. Thus, a STEM internship has the ability to a) expose them to new academic and career options, b) provide them with the self-efficacy and confidence to succeed, and c) integrate them into the STEM community at a critical time in their undergraduate studies. Our findings reveal that – years after completing the program – students who received quality mentoring and support retained positive memories and associations regarding their experience.

## Recommendations

### Partnerships with community colleges

Many community colleges are disconnected from the STEM professional community, have less access to engagement in internships and research experiences for their students, and promote employment over professional development opportunities that would support the development of STEM identity.^50,139^ Programs that involve partnerships between community colleges and other institutions can be an effective way to support student success during “the transitions from one part of their career pathway to another.”^165^ Aligned with the call to action by Hampton-Marcell and colleagues^2^ to support Black students through partnerships between schools and DOE national laboratories, we recommend that laboratories establish strategic partnerships with community colleges to provide students with “early exposure” research opportunities. Considering previous findings about community college student preference for academic/career opportunities within a “comfortable distance,” we recommend that DOE national laboratories engage in outreach efforts that include community colleges in the surrounding geographic area.^140–143^ Additionally, DOE national laboratories can communicate with program alumni to share information about future opportunities to enter the DOE workforce and engage in outreach efforts that include schools in the surrounding geographic area, to provide opportunities to both students and faculty at community colleges.^166,167^

### Applicant evaluation and selection

Broadening the scope of possible ways to evaluate program applicants is one way to address the disparities in resources across different student populations and ensure that a diverse new generation of STEM professionals is trained and supported to succeed from the undergraduate level and beyond. Programs should also consider the bias that may be present in their eligibility requirements, application structure, selection criteria, and/or recommendation letters against those student populations with less access to STEM careers. For example, an applicant’s GPA may not be reflective of their disposition and interest in working on research/technical projects.^17^ Programs could reduce bias by the use of a standardized recommendation letter, which would produce “a similar description for each student” applicant.^149^ Those involved in reviewing applications could consider the potential impact such an opportunity might have for a student with limited access to those opportunities. The Level Playing Field Institute in Oakland, California and the Biology Scholars Program at the University of California, Berkeley consider factors such as “distance traveled.” Rather than previous achievements, *distance traveled* examines an applicant’s trajectory, including the resources and support they had access to and what hurdles they have overcome to arrive at their current academic/career stage.^168–171^ Similarly, McDevitt and colleagues^150^ suggest a two-step approach, in which program directors first review and narrow the applicant pool based on project needs, program goals, etc., and then mentors select students from this pool based on skills and their “potential to gain additional value” from the program.

### Practices in support of community college students

For students and early career professionals, positive and supportive mentor-mentee relationships in a particular professional environment contribute to increased desire to remain in a similar career pathway.^152^ Creating a positive and supportive working environment is beneficial to all parties in the short-term, and can have long-term impacts on students, as well. Mentors will best serve all students by being aware of the possible ways in which background, culture, and identity can impact students’ academic/career experiences and perspectives. Unlike many of the resources needed to offer a professional development opportunity to students, kindness is free and readily available for all members of the STEM professional community to give to others. Although we often associate professional development opportunities with productivity and career advancement, mentors whose practices include kindness, attention, and trust can have many positive impacts on their mentees years into the future.

Mentors, counselors, and staff can expose and challenge negative stereotypes about community colleges, to support students’ pride in their educational pathways and identities and increase the likelihood that they will persist and complete their studies.^42,147,172^ We recommend that those individuals involved in the recruitment, training, and education of community college and transfer students learn about these issues and take active steps to empower these students.

## Conclusions and Future work

Based on our in-depth communication with individuals who were interested in STEM disciplines as community college students, we understand some of the reasons why they initially held low expectations of being successful in the STEM workforce: they did not understand what science/engineering entailed, and/or they did not have the support to pursue their interests in these disciplines. We also learned that, to retain students in STEM career pathways, it is not enough to recruit them into technical or research experiences. The ways in which STEM professionals, program staff, guest speakers, and other members of the community interact with students are critical to their professional development and perception that they are capable of completing STEM degrees and entering the STEM workforce.

Our study made use of the existing SCCT model, which helped us to interpret our findings in the context of previous scholarship about STEM research and technical experiences. Indeed, CCI at LBNL serves as a SCCT-aligned learning experience for community college students, influencing their academic/career interests, goals, and actions. We propose several additions to the SCCT model, to better reflect the supports and barriers to STEM persistence for community college students.

Many studies about internships provide insufficient detail about program structure, and fail to connect internship characteristics with student outcomes.^57,173^ It is not enough to report the goals of a program; we urge more scholars to contribute to knowledge about STEM professional development for community college STEM majors by publishing studies that clearly describe program elements and connect these to participant outcomes. The academic pathways and learning environments of community college students can be very different than those of students attending baccalaureate-granting institutions, so interventions, assessment strategies, and research studies should be developed with partners who possess expertise about this unique student population.^139,174–176^ Additionally, we encourage faculty and scholars with ties to community colleges to be involved in studies about the experiences of community college students, interventions beneficial to them, and the development of new approaches to support STEM learning and workforce development. Similarly, studies about the STEM community, academic/career pathways, and psychosocial elements of learning environments will be better able to support inclusion, high-quality programming, and retention in the STEM workforce when designed in partnership with educators and social scientists.^24,177^ Considering the rare mention of programs at DOE national laboratories in the research literature, we advocate for collaborations between STEM professionals and those with training and expertise in educational research and social sciences to study this topic. Scholarship in this area has the potential to influence policy, funding, and the adoption of new ideas for impactful and inclusive learning environments.

## Acknowledgements

We are grateful to Laura Almaguer, Christine Beavers, Christopher Chen, Jerica Duey, Katie Dunne, Brett Heischmidt, Chris Martinez, Melissa Meikle, and Daniel Ramos for supporting the research team during our early development and piloting of the survey and interview protocols. Dianna Bolt, Sean Burns, Chris Byrne, Christel Cantlin, Joseph Crippen, Nakeiah Harrell, Lady Idos, and Christina Teller assisted with records and information. Finally, thanks to Sean Burns, Krista Cortes, Amanda Dillon, Ali Esmaili, Max Helix, Megan Hochstrasser, Kelsey Miller, Rey Morales, Erin Murphy-Graham, Sinéad Griffin, Lloyd Goldwasser, Kris Gutiérrez, Jennifer King Chen, Ingrid Ockert, Miguel Órdenes, Colette Patt, Christopher Payne, Laura Pryor, Michael Ranney, Renee’ Schwartz, Angy Stacy, Elisa Stone, Alice Taylor, Matt Traxler, Karen Villegas, Michelle Wilkerson, Brieanna Wright, Rachel Woods-Robinson, Sandra Zuñiga-Ruiz, and the Spring 2022 EDUC 209 class at the University of California–Berkeley for their feedback. L.E.C. would like to acknowledge the role that Rey Morales, Eduardo Cervantes, and Darlene Del Carmen played in “nudging” students (like her) at Gavilan College to apply to professional development opportunities.

## Author Contributions

**Conceptualization:** Laleh E. Coté, Colette L. Flood, Anne M. Baranger

**Data curation:** Laleh E. Coté, Seth Van Doren, Astrid N. Zamora, Julio Jaramillo Salcido, Esther W. Law, Gabriel Otero Munoz

**Formal Analysis:** Laleh E. Coté, Seth Van Doren, Astrid N. Zamora, Julio Jaramillo Salcido, Esther W. Law, Gabriel Otero Munoz, Aparna Manocha

**Funding acquisition:** Laleh E. Coté, Colette L. Flood

**Investigation:** Laleh E. Coté, Seth Van Doren, Astrid N. Zamora, Julio Jaramillo Salcido, Esther W. Law, Gabriel Otero Munoz, Aparna Manocha

**Methodology:** Laleh E. Coté, Anne M. Baranger

**Project administration:** Laleh E. Coté

**Supervision:** Anne M. Baranger

**Validation:** Laleh E. Coté, Seth Van Doren, Anne M. Baranger

**Visualization:** Laleh E. Coté, Seth Van Doren, Julio Jaramillo Salcido, Aparna Manocha, Anne M. Baranger

**Writing – original draft:** Laleh E. Coté

**Writing – review & editing:** Laleh E. Coté, Seth Van Doren, Astrid N. Zamora, Julio Jaramillo Salcido, Colette L. Flood, Anne M. Baranger

## Data Availability

The data collected contains information that could compromise the privacy of research participants, so the full dataset generated for the current study is not publicly available. However, a selection of data that support the findings of this paper will be available from the corresponding author upon reasonable request.

