## Supplemental Materials for "“When I talk about it, my eyes light up!” Impacts of a national laboratory internship on community college student success"

**Short title:** National laboratory internship and community college student success

**Authors:**
Laleh E. Coté*, Lawrence Berkeley National Laboratory, https://orcid.org/0000-0002-6232-7573
Seth Van Doren, University of California, Irvine, https://orcid.org/0000-0003-0674-277X
Astrid N. Zamora, Stanford University, https://orcid.org/0000-0001-8187-3492
Julio Jaramillo Salcido, Lawrence Berkeley National Laboratory, https://orcid.org/0000-0002-4113-5345
Esther W. Law, Milliman, Inc.
Gabriel Otero Munoz, Indiana University Bloomington, https://orcid.org/0000-0002-1444-8020
Aparna Manocha, University of California, San Francisco, https://orcid.org/0000-0001-7824-9971
Colette L. Flood, Lawrence Berkeley National Laboratory, https://orcid.org/0000-0002-8674-0872
Anne M. Baranger, University of California, Berkeley, https://orcid.org/0000-0002-1973-4632

***Corresponding author:**
Laleh E. Coté, Ph.D.
Workforce Development & Education
Lawrence Berkeley National Laboratory
1 Cyclotron Road Mailstop 50A1148
Berkeley, CA 94720


**Supporting Information**

**S1 Figure. *Community College Internship (CCI) Alumni Survey*, developed for use with CCI alumni.**

| Questions |
| --- |

First Name

Last Name

If applicable, please provide your previous name(s) used at Berkeley Lab (during your participation in the CCI program).

Contact email address(es)

Please select the term in which you first participated in the CCI program at Berkeley Lab:

If you completed a second CCI term at Berkeley Lab, please select the term:

If you participated in another internship program at Berkeley Lab (e.g., 1st term of SULI), please select the term:

If you participated in additional terms of another internship program at Berkeley Lab (e.g., 2nd term of SULI), please select the term:

Please explain why you initially decided to apply to the CCI program at Berkeley Lab.

If you completed more than one term in any of our internship programs, how did you feel that participating in an additional term would benefit you?

Did you graduate from community college? (Select all that apply)

- ☐ Yes, I obtained an A.A. or A.S. degree (STEM field).
- ☐ No, I'm currently attending community college (majoring in a STEM field).
- ☐ Yes, I obtained an A.A. or A.S. degree (non-STEM field).
- ☐ No, I'm currently attending community college (majoring in a non-STEM field).
- ☐ No, I transferred to a 4-year university without obtaining an A.A. or A.S. degree.
- ☐ No.
- ☐ I decline to state.

Did you attend a 4-year university? (Select all that apply)

- ☐ Yes, I obtained a B.A. or B.S. degree (STEM field).
- ☐ Yes, I attended a 4-year university (majoring in a STEM field), but did not graduate.
- ☐ Yes, I'm currently attending a 4-year university (majoring in a STEM field).
- ☐ Yes, I obtained a B.A. or B.S. degree (non-STEM field).
- ☐ Yes, I attended a 4-year university (majoring in a non-STEM field), but did not graduate.
- ☐ Yes, I'm currently attending a 4-year university (majoring in a non-STEM field).
- ☐ No, I'm currently attending community college.
- ☐ No.

- ☐ I decline to state.

Did you attend graduate school? (Select all that apply)

- ☐ Yes, I obtained an M.A. or M.S. degree (STEM field).
- ☐ Yes, I obtained a Ph.D. degree (STEM field).
- ☐ Yes, I attended graduate school (STEM field), but did not graduate.
- ☐ Yes, I'm currently attending graduate school (STEM field), with the goal of obtaining an M.A. or M.S. degree.
- ☐ Yes, I'm currently attending graduate school (STEM field), with the goal of obtaining a Ph.D. degree.
- ☐ Yes, I obtained an M.A. or M.S. degree (non-STEM field).
- ☐ Yes, I obtained a Ph.D. degree (non-STEM field).
- ☐ Yes, I attended graduate school (non-STEM field), but did not graduate.
- ☐ Yes, I'm currently attending graduate school (non-STEM field), with the goal of obtaining an M.A. or M.S. degree.
- ☐ Yes, I'm currently attending graduate school (non-STEM field), with the goal of obtaining a Ph.D. degree.
- ☐ Yes, I am currently attending graduate school or a professional school not represented by the choices given (e.g., medical school, law school, business school).
- ☐ Yes, I completed graduate-level studies or a professional school not represented by the choices given (e.g., medical school, law school, business school).
- ☐ No, I'm currently attending community college or a 4-year university.
- ☐ No.
- ☐ I decline to state.

Briefly describe your recent educational, professional, extracurricular, and/or personal activities.

Do you feel that your needs as a student were addressed and/or met by participating in the CCI program at Berkeley Lab? Why, or why not?

Briefly describe your recent educational, professional, extracurricular, and/or personal activities.

Briefly describe your "dream job", and why you would like to engage in that type of work.

Please specify in which of the following fields you have been employed, or have professional experience.

In the future, in which of the following fields are you interested in working? (Select all that apply)

- ☐ Academia
- ☐ Industry
- ☐ Government
- ☐ Research
- ☐ STEM policy
- ☐ STEM field, which requires use and/or knowledge of technical skills
- ☐ STEM field, which does not require use and/or knowledge of technical skills
- ☐ U.S. Department of Energy national laboratory
- ☐ U.S. Department of Energy facility (not a national laboratory)
- ☐ STEM education and/or outreach

- ☐ Science media and/or communication
- ☐ Non-STEM field, which requires use and/or knowledge or technical skills
- ☐ Non-STEM field
- ☐ None of these

Since participating in the CCI program, in what capacity have you worked at Berkeley Lab? (Select all that apply)

- ☐ Intern in another program
- ☐ Undergraduate researcher
- ☐ Post-baccalaureate researcher
- ☐ Research associate or research assistant
- ☐ Graduate student researcher
- ☐ Post-doctoral researcher
- ☐ Employee
- ☐ Technical staff
- ☐ None of these

In the future, in which ways are you interested in working at Berkeley Lab? (Select all that apply)

- ☐ Intern in another program
- ☐ Undergraduate researcher
- ☐ Post-baccalaureate researcher
- ☐ Research associate or research assistant
- ☐ Graduate student researcher
- ☐ Post-doctoral researcher
- ☐ Employee
- ☐ Technical staff
- ☐ None of these

Since participating in the CCI program, in what capacity have you worked at any U.S. Department of Energy national laboratory or facility? (Select all that apply)

- ☐ Intern in another program
- ☐ Undergraduate researcher
- ☐ Post-baccalaureate researcher
- ☐ Research associate or research assistant
- ☐ Graduate student researcher
- ☐ Post-doctoral researcher
- ☐ Employee
- ☐ Technical staff
- ☐ None of these

What factors do you believe are important for having a successful research experience at Berkeley Lab?

In your opinion, what makes the undergraduate research experiences at Berkeley Lab different or unique from other internships?

Are you still in touch with any of the members of your Mentor Group? If so, in what capacity?

As a result of your work during the CCI program at Berkeley Lab, please list any related project outcomes, such as publications, presentations, additional collaborations, employment opportunities, etc.

How did your experiences at Berkeley Lab influence your academic or career plans?

*If possible, please elaborate on how these experiences did or did not change your perspectives about what it is like to work in research and/or science.*

Which activities or experiences from the CCI program made the biggest impact on your career, and why?

Please describe (or give examples) of ways in which you feel that engaging in undergraduate research experiences at Berkeley Lab prepared you to solve problems on other projects, or at other organizations.

What were the most valuable aspects of your experience in the CCI program, and why?

What are ways that the program could be improved to better support community college students?

Please share any memorable stories from your experiences in the CCI program.

*These may be experiences which influenced your career goals, had a personal impact on you, changed your perspectives about working in a research-based environment, left you feeling empowered or frustrated, or challenged you in some way. Anything that comes to mind is fair game, and you are encouraged to share as much or as little as you feel comfortable with. Please note that any personally identifiable information about you or others (including names of individuals and group names) will be made anonymous.*

Please share any times when you, as an undergraduate, felt like a scientist.
*(Or, researcher, biologist, chemist, physicist, mathematician, computer scientist, engineer, etc.)*

Please share any other comments you might have about your experiences in the CCI program or working at Berkeley Lab.

What do you think would be an effective way to encourage community college students to apply to the CCI program at Berkeley Lab, or any undergraduate research experience?

---

**S2 Figure. Semi-structured interview protocol, developed for use with CCI alumni.**

| Questions |
| --- |

**Section A.**
**Tell me a bit about you. What is your name, where are you from, and where do you work or go to school?**

Why did you attend college?

What did you major in?

How did you become interested in this topic?

Can you describe what it felt like to be an undergraduate student, just beginning to study science or engineering? (use their field of study)

- How confident were you in your general research or technical skills?
- How confident were you in your ability to succeed in graduate school?
- How confident were you in your ability to succeed in the STEM workforce?

What words would you use to describe your identity?

How does <your field of study> fit in there?

Before becoming involved in research as an undergrad, did you ever feel like a scientist or engineer? (use term they chose)

What are you good at, that makes you well-suited for <their field of study>?

In <their field of study>, who are the people you identify with?

**Section B.**
**Let's talk about the CCI program at Berkeley Lab now.**

What happened that made you want to apply to the program in the first place?

Can you briefly describe the type of research you worked on?

What was the benefit of working on-site at Berkeley Lab, instead of collaborating with your team remotely?

Can you share any stories with me about times when you felt successful, as an intern?

What about times when you might have felt unsuccessful?

During the program, how much autonomy did you have as an intern?

How much did you collaborate with others on your CCI project?

When you were an intern, what kinds of conversations would you typically have with your mentors?

In what ways do you feel that participation in this program impacted you personally?

Were there times during the program when you really felt like a scientist or engineer? (use term they chose)

After you completed the CCI program:

- How confident were you in your general research or technical skills?
- How confident were you in your ability to succeed in graduate school?
- How confident were you in your ability to succeed in the STEM workforce?

**Section C.**
**We're almost done! During the next few questions, we will discuss your future.**

What are your future academic or career goals?

Thinking now about everything we've discussed, what aspects of your college experience really impacted how you might go about achieving those goals?

Let's pretend for a moment that you have a sibling who is 5-6 years younger than you. Inspired by your career path, your sibling enrolls in the same community college you attended, and declares the same exact major. What advice would you give them, or what strategies would you recommend to them, to support their success in this field?

Can you share any other experiences that you felt were important, that we haven't already discussed?

---

**S1 Table. Self-reported characteristics of CCI alumni interviewed for this study.**

| Characteristic | | interview subjects | |
|---|---|---|---|
| | | n | % |
| STEM field of study | | | |
| | Civil and/or mechanical engineering | 4 | 33 |
| | Physics and/or mathematics | 3 | 25 |
| | Chemistry | 2 | 17 |
| | Biology | 2 | 17 |
| | Environmental Science | 1 | 8 |
| School location | | | |
| | Attended a California community college | 10 | 83 |
| | Attended a community college outside of California | | |
| Academic achievement | | | |
| | Has a B.A./B.S. in STEM | 11 | 92 |
| | Has an M.A./M.S. in STEM | 5 | 42 |
| | Has a Ph.D. in STEM | 2 | 17 |
| | Studied STEM after obtaining a non-STEM B.A./B.S.[a] | 2 | 17 |
| | Has an advanced degree in health (Ph.D., M.D., D.D.S.) | 1 | 8 |
| Current academic or professional activity | | | |
| | Studying/working in a STEM field | 11 | 92 |
| | Working at a DOE national laboratory | 3 | 25 |
| | Attending graduate school for a STEM Ph.D. | 3 | 25 |

| | | | |
|---|---|---|---|
| | Studying/working in a health field | 1 | 8 |
| STEM perspectives | | | |
| | First in family to study science | 8 | 67 |
| | Perceptions and experiences in STEM impacted by background, culture, and/or identity | 6 | 50 |
| | Passionate about STEM education and outreach | 6 | 50 |
| | "Always" liked science | 5 | 42 |
| | Believes STEM is a pathway to upward mobility | 4 | 33 |
| | Interested in how philosophy and science intersect | 3 | 25 |
| | Became interested in STEM in high school | 2 | 17 |
| Other self-identified characteristics | | | |
| | From a low-income family | 5 | 42 |
| | Non-traditional age (during undergraduate studies) | 4 | 33 |
| | First-generation to college | 4 | 33 |
| | Working-class | 3 | 25 |
| | Parent | 3 | 25 |
| | From a rural community | 2 | 17 |
| | Immigrant | 1 | 8 |
| | Child of immigrants | 1 | 8 |

The characteristics of each individual are described by more than one category. [a] These individuals obtained a degree in a non-STEM subject and entered the non-STEM workforce before re-entering school to take STEM coursework at a community college.

**S2 Table. Interview responses from CCI alumni about confidence in being successful in the STEM workforce.**

| Before CCI, how confident were you in your ability to succeed in the STEM workforce? | After CCI, how confident were you in your ability to succeed in the STEM workforce? |
|---|---|
| "... in terms of getting into a science career, I felt a low sense of confidence before CCI." | "I felt that was much more of a possibility than ever before." |
| "I wasn't thinking about it, it was more like, 'I'm going to college and then I'm getting a job.' I didn't think about what." | "I think so, because now I had an idea of something I wanted to do in the future. Since I had a crack at it, it was … something I [could] see myself doing and feel confident doing." |
| "Pretty low. Yeah, I didn't have much confidence. I thought I'd just do physics as a hobby." | "Again, I wouldn't say extremely confident, but from low to mildly confident." |
| "Yeah, yeah, I think I was. That's why I wanted to have an engineering degree. I feel like it was very straight-forward. I know how to do this skill, and there's a job I could fulfill that requires that skill." | "... it wasn't like I hadn't had experience in the job workforce before. Talking to well-educated academics, people of that different kind of caliber, … I think that was more of an experience for me." |
| "Not particularly." | "More confident. It [showed] me a different aspect of the workforce where, I think, I felt I excelled more." |

During interviews, we asked the following two questions: As an undergraduate (before CCI), how confident were you in your general research or technical skills? After you completed the CCI program, how confident were you in your general research skills? These are a selection of the responses we received from CCI alumni, which are representative of the individuals we interviewed (n=12). Each row contains two quotes, and these are both from the same individual.

**S3 Table. Interview responses from CCI alumni about confidence in their general research or technical skills.**

| Before CCI, how confident were you in your general research or technical skills? | After CCI, how confident were you in your general research or technical skills? |
|---|---|
| “Not very. When I first started community college the thought of doing research was ... I didn’t even understand that research was something you do in an academic setting.” | “I felt stronger ... I don’t know if I considered myself like a great researcher, but I remember at the Lab we did, there was this whole component with CCI where we were doing … some sort of report ... and I came up with this whole research thesis ...” |
| “No. I was just like, ‘books, study, books, study, books, study.’ ” | “After I got done I was like, ‘Oh, yeah, I can definitely do something else research-wise,’ and now I can actually probably start my data collection and my analysis with whatever tools I needed. Yeah, I definitely have some understanding of how I would set up to study something.” |
| "I couldn’t grasp what scientists do. I could understand that chemists wear lab coats and do titrations. So, basically, I had no understanding." | “I felt you know, I could do research. Again, I don’t remember one moment of clarity, like, ‘I know this!’ But, it was a gradual process. It was high, I’d say.” |
| “I had no experience with general research.” | “[The] reality was, that I wasn’t that confident in my lab skills, but I really felt like I’d come a long way in putting together that research paper. I think just the process of writing a paper and doing a poster presentation and going from researching this background of this field and connecting that with my own research, ... it felt like I had gained a lot. For sure.” |
| “At the time, I didn’t know what research was.” | “Very confident. Now I knew more of what that entails ... when it comes to doing research in biology and that kind of field, when it comes to wet lab stuff, way more confident.” |

During interviews, we asked the following two questions: As an undergraduate (before CCI), how confident were you in your general research or technical skills? After you completed the CCI program, how confident were you in your general research or technical skills? These are a selection of the responses we received from CCI alumni, which are representative of the individuals we interviewed (n=12). Each row contains two quotes that are from the same individual.

**S4 Table. Interview responses from CCI alumni about confidence in being successful in graduate school.**

| Before CCI, how confident were you in your ability to succeed in grad school? | After CCI, how confident were you in your ability to succeed in grad school? |
|---|---|
| "I didn't put thought into that. It was, I knew college was a thing I should do, but I didn't think past that." | "At that point, I was actually thinking about that more. I was confident enough that was what I wanted to do. I could do it." |
| "... it was on my radar, but I didn't really. I needed to know exactly what I was going to end up focusing on in civil [engineering]. … I don't think I had thought of research as something I wanted to focus on." | "Well yeah, [CCI] was like grad school 101. That was like, 'okay, this is a little tiny version of what you have to do in grad school.' So, yeah, for sure, that was … I don't know how much closer you can get in an internship with how he worked with us, you know." |
| "Oh yeah, no. I didn't think I could handle community college! Like, grad school was something other people did. No." | "Definitely was thinking about it. I felt pretty confident that I could get to grad school. I had a better understanding of what that entailed, and what that looked like. Like I said before, we were joking about how we were all going to get our PhDs. Now that was an option." |
| "I might've had an inflated sense of confidence because I had no idea what it would be like. I was so out of touch, I didn't really know what a PhD was." | "I felt excited and optimistic about my ability to [succeed] in grad school afterwards." |
| "No, I thought that would be a dead end. Yeah, so I, in the back of my mind, I thought I'd do engineering, so at least I could get a job ... That sounded the most financially viable path for me at the time." | "After research, after [the] CCI program, I said, 'absolutely I will go for graduate school.' I think the feeling that I could do research, because there was always a lack of confidence. I thought I was average, and how could I contribute to research? But, going in and doing research, I felt I could make a contribution. That all fell nicely in front [of] me." |

During interviews, we asked the following two questions: As an undergraduate (before CCI), how confident were you in your ability to succeed in graduate school? After you completed the CCI program, how confident were you in your ability to succeed in graduate school? These are a selection of the responses we received from CCI alumni, which are representative of the individuals we interviewed (n=12). Each row contains two quotes that are from the same individual.

**S5 Table. Survey responses from CCI alumni about their "dream jobs."**

| Describe your "dream job" |
| --- |
| "I want to be a technical manager. I would have to do both technical and managerial work. Also, it would be at a company that cares about having a good workplace culture and investing in its employees." |
| "I would ideally like to be a research scientist at a DOE National Lab … studying nuclear reactions as they pertain to nuclear astrophysics. I am interested in understanding the origins of the elements in our universe, which many government and university labs are working towards." |
| "I want to be a research scientist working in a collaboration on a large experiment. I enjoy working with diverse groups of people and I would enjoy … choosing what direction I want my research to go in." |
| "I would like to do the mechanical work that can best maximize efficiency [for] major utilities … figuring out what would be the best equipment to transport water from treatment plants, or designing the best route for electricity to travel with the least amount of power loss." |
| "I would like to work as a pharmaceutical liaison acting as a medical and scientific expert engaged in driving key initiatives in research, publications, medical education and field intelligence ... collaborating with researchers and physicians to develop new life saving drugs and treatments." |
| "My dream job was to be a renewable energy scientist at [DOE national lab] or similar … because I am deeply passionate about the intellectual challenges and excitement of working with cutting edge people." |
| "My dream job would be to be a design engineer or process engineer for companies such as [for-profit company] or [for-profit company] … because the work of an engineer is directly related to the advances we see in this world today. It is always exciting to say that 'I have been a part of this great invention.' " |
| "I'd like to work in computational research focusing on the ocean and atmosphere. Computer programming keeps me excited to solve problems every day, and applying it to natural science keeps me interested and passionate about my work on longer time scales." |

In the survey we asked CCI alumni to respond to the following prompt: Briefly describe your "dream job," and why you would like to engage in that type of work. These are a representative selection of the responses we received.

**S6 Table. Coding categories, codes, and sub-codes applied to survey and interview data.**

| Category from SCCT model | Coding category | Generated codes | Generated sub-codes |
|---|---|---|---|
| Personal inputs | Personal inputs | Gender | |
| | | Race/ethnicity | |
| | | First-generation to college | |
| Background contextual influences | Background contextual influences | Pre-program social supports and barriers | Attitudes toward community college |
| | | | Support from family and friends (of STEM academic/career goals) |
| Proximal contextual influences | Proximal contextual influences | Pre-program social supports and barriers | Support from people associated with the community college |
| | | | Learning experiences available to community college students |
| | | Social supports and barriers | Support from people associated with the learning experience |
| | | | Established network associated with the learning experience |
| | | | Kindness from people associated with the learning experience |
| | | | Mentoring received during the learning experience |
| Learning experiences | Learning experiences | Pre-program learning experiences (Courses, clubs, other opportunities to learn about STEM) | |
| | | Learning experience (Community College Internship at LBNL) | |
| | Skill development | Pre-program STEM skills | |
| | | STEM skills | |
| | Knowledge about STEM careers | Knowledge about STEM careers | |

| | | | |
|---|---|---|---|
| Self-efficacy | Self-efficacy, confidence, STEM identity | Self-efficacy | |
| | | Confidence | |
| | | Feeling like a scientist or engineer (STEM identity) | |
| Outcome expectations | Outcome expectations | Pre-program STEM outcome expectations | Expectations of admittance into the learning experience (CCI) |
| | | | Expectations of success in a STEM career |
| | | | Expectations about working in research |
| | | | Expectations of graduating from a baccalaureate granting institution |
| | | | Expectations of attending graduate school |
| | | STEM outcome expectations | Expectations of success in a STEM career |
| | | | Expectations about working in research |
| | | | Expectations of graduating from a baccalaureate granting institution |
| | | | Expectations of attending graduate school |
| Interests | Academic and career interests | Pre-program STEM interests | |
| | | STEM interests | Interest in a specific research or STEM field/topic |
| Choice Goals | Academic and career goals | Academic and career goals | |
| Choice Actions | Actions | Academic and career actions | |
| Persistence | Persistence | Persistence in STEM | |
